# Contextual inference through flexible integration of environmental features and behavioural outcomes

**DOI:** 10.1101/2025.05.28.656607

**Authors:** Jessica Passlack, Andrew F MacAskill

## Abstract

The ability to use the context we are in to flexibly adjust our decision-making is vital for navigating a complex world. To do this, the brain must i) use environmental features and behavioural outcomes to distinguish between different, often hidden contexts; and ii) learn how to use these inferred contexts to guide behaviour. However, how these two interacting processes can be performed simultaneously remains unclear. Within the brain it is thought that interaction between the prefrontal cortex (PFC) and hippocampus (HPC) supports contextual inference. We show that models of the HPC using environmental features readily support context-specific behaviour, but struggle to differentiate ambiguous contexts during learning. In contrast, models of the PFC using behavioural outcomes can stably differentiate contexts during periods of learning, but struggle to guide context-specific behaviour. Supporting feature-based with outcome-based strategies during learning overcomes the limitations of both approaches, allowing for the formation of distinct contextual representations that support contextual inference. Moreover, agents using this joint approach reproduce both behavioural- and cellular-level phenomena associated with the interaction between PFC and HPC. Together, these results provide insight into how the brain uses contextual information to guide flexible behaviour.

## INTRODUCTION

Humans and animals have a remarkable ability to learn and recall multiple optimal behaviours dependent on the current context [1, 2, 3, 4, 5]. While such contexts are often distinct environments that are easily distinguishable from one another, distinct contexts are also commonly found within the same environment where they can be differentiated by the presence of distinct features (such as smells, objects or sounds), or the distinct distribution of outcomes (such as rewards). This ability to identify distinct contexts and differentiate between them - often termed contextual inference - is crucial, as different behaviours are needed to achieve the optimal outcome in each context. Importantly, while the brain can effortlessly perform contextual inference, how this is achieved remains unclear, apart from knowing that it requires complex interplay between multiple brain regions, in particular the prefrontal cortex and hippocampus [6, 7]. Similarly, computational approaches are still limited in their ability to simultaneously learn to identify contexts and drive context-dependent behaviour [8, 9].

Computationally, two main approaches have had success with learning to perform contextual inference. One approach is to use recurrence to maintain information related to the hidden context over time in the form of recurrent neural networks (RNNs) [10, 11, 12, 13]. The other approach uses approximate Bayesian inference over model-based representations that learn explicitly how observations are linked to each hidden context [14, 15, 1, 16, 17]. However, despite computational approaches being able to successfully maintain multiple context-dependent representations [12, 16, 18], during learning these approaches often struggle to identify the correct context upon which to learn, especially during more complex tasks. Specifically, models struggle to learn when the signal-to-noise ratio (SNR) of context-dependent to non-context-dependent observations decreases [13, 10, 19, 20]. This inability to cope with low contextual SNR is a major limitation to applying these techniques successfully to complex environments as, given the noise and uncertainty of the world, often proportionately small changes signal a change in the current optimal behaviour. Recent evidence shows that this issue can be overcome by using externally provided context signals during learning, however how such supporting context signals can be generated and how they support learning remains unclear [21, 22, 23, 24].

Interestingly, in the brain, regions that support contextual inference also seem to rely on support from other brain regions specifically during learning. In particular, prefrontal cortex (PFC) input to the hippocampus (HPC) is important during learning [25, 26, 27, 28]. The HPC is thought to learn context-dependent, detail-oriented maps of features like sounds, location, time, and odours that allow for the prediction of upcoming features based on currently observed features [29, 30, 31, 32, 33, 1], similar to RNNs [34] and Bayesian model-based representations [35, 36]. In contrast, the PFC is often proposed to classify different contexts based on their distinct rules. More specifically, PFC is thought to differentiate between contexts based on the distinct outcomes that result from following each rule in each context [37, 28, 38, 39, 6, 39, 40, 2]. As a result, within the brain it seems that outcome-based approaches in PFC may support the generation of a context signal that can support the learning of more complex, feature-based approaches in HPC.

Computationally, both feature- and outcome-based approaches can predict the current context based on their experience within the environment. An example of using features to infer the current context is adjusting the amount of flour you add to bread dough on the fly depending on the appearance and texture of the dough. However, if it is your first time making that specific bread and you do not know what the dough should look like, you can instead use the outcome of whether the resulting loaf is too dry or too wet to adjust the amount of flour you add next time. As is evident in this example, feature- and outcome-based approaches trade-off in their ability to predict contextual changes within a trial and in the complexity of the representations they use. Feature-based approaches learn complex, detailed representations of the environment, whereas outcome-based approaches focus on learning only about one parameter - outcomes associated with behaviour.

The greatly reduced complexity of learnt representations in outcome-based approaches compared to feature-based approaches allows outcome-based approaches to rapidly learn to distinguish different contexts [41, 42, 43]. However, by design outcome-based approaches can only react to changes in behavioural outcomes once they have been experienced. In contrast, although they are more complex, feature-based approaches are able to predict changes in outcome in advance of them being experienced, by identifying changes in the environment using the detailed representations they have created. On this basis, in this study we hypothesised that during learning - especially where distinct contexts have increasingly overlapping features - the initial differentiation of contexts by feature-based models may benefit from the addition of a stable ‘teacher’ in the form of an outcome-based algorithm.

Therefore in this study we first characterise the performance of agents utilising each of these strategies in a simple, context-dependent task. We then go on to probe how these agents cope with tasks with decreasing contextual SNR, and ask to what extent combining both strategies mitigates these impairments. Finally, we test the generalisability of our findings by investigating how these algorithms perform on a series of common behavioural tasks used in the neuroscience community.

## RESULTS

Our main goals were to investigate i) the mechanism underlying deficits in learning to perform contextual inference that occur as contextual SNR decreases and ii) whether such deficits could be alleviated by incorporating outcome-based inference during learning. To test the performance of outcome- and feature-based approaches, we trained and probed the learning of these algorithms on a simple cued T-maze paradigm often used in both neuroscience and computational experiments investigating contextual inference (**Figure 1A**). In this task a distinct cue at the beginning of a T-shaped maze indicates whether the left or right arm is rewarded [10, 23, 44, 45, 46]. This choice of task facilitated comparison of the performance and workings of our algorithms with neuroscience data (see also **Figure 7** for comparison with other common neuroscience tasks).

**Figure 1.**
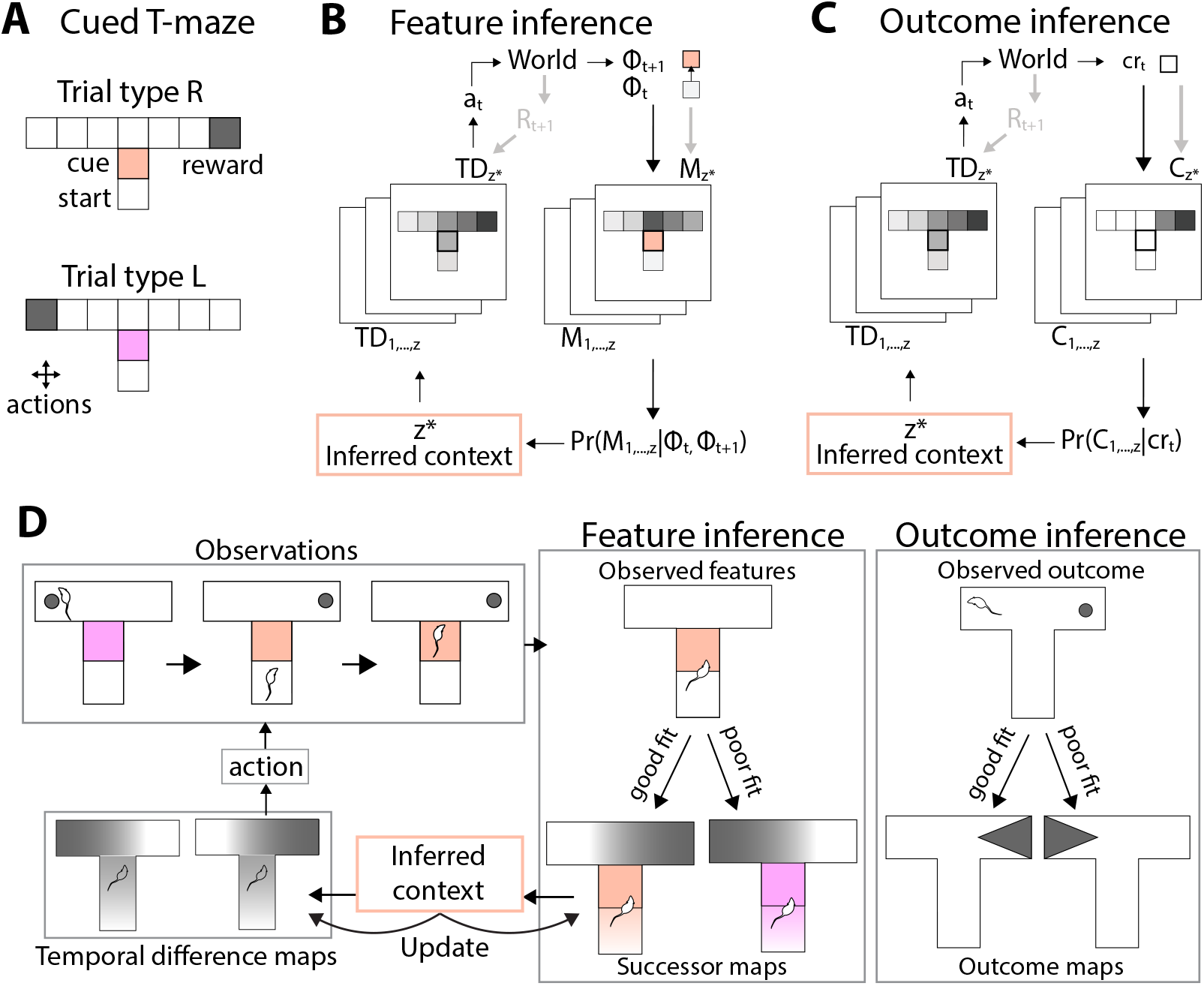
Design of feature and outcome inference models. (A) Setup of the cued T-maze task, showing different location and cue-based features, (B) Algorithm underlying feature inference showing how observed feature transitions *ϕ*_*t*_ to *ϕ*_*t*+1_ are compared with learnt successor feature maps *M* using Bayesian inference to determine which context *z*^∗^ is currently most likely, followed by using the corresponding temporal difference map *T D* to choose the current best action *a*_*t*_, (C) Algorithm underlying outcome inference showing how observed outcomes *cr*_*t*_ are compared with learnt convolved reward maps *C* using Bayesian inference to determine which context is currently most likely, (D) Schematic of how the feature and outcome inference models react to a change in trial type, showing selection of the relevant map for feature inference following the cue and for outcome inference following the lack of reward, and how the inferred context allows for action selection using separate temporal difference maps.

We modelled the T-maze task by representing each arm of the maze as 3 discrete locations - leading to 9 total discrete locations. We then represented the cue as an additional discrete feature at one single location in the starting arm (**Figure 1A**). To perform the task, an agent must choose actions to transition between discrete locations. After every step, the agent observes its current location and whether there is a cue present. Each ‘attempt’ ends when the agent reaches the end of the left or right arm, after which the agent is teleported back to the starting state. The agent has an unlimited number of attempts to make the correct choice and once the correct choice is made it moves into the next trial.

We focused on a Bayesian model-based approach, to allow us to easily probe how learnt representations support behaviour. Specifically, to model feature-based strategies, we used a Bayesian successor feature algorithm, which we refer to as ‘feature inference’ (FI). This model infers and predicts optimal behaviour based on differing features of the environment using context-specific successor representations (SRs) [14]. To model outcome-based strategies, and to investigate whether these can be used to support learning of feature inference, we used a similar model-based Bayesian approach, which we refer to as ‘outcome inference’ (OI). In contrast to feature inference, this model learns context-specific diffuse state-outcome maps to identify contexts [42]. As outcome inference focuses its attention solely on state-outcomes, which define the optimal behaviour, we hypothesised that it would be well suited to support initial learning of context-specific SRs for predictive inference.

To solve this task, both FI and OI learn a set of context-dependent maps of the world, which they use to identify the current context. FI learns a set of SRs *M* and OI learns a set of convolved reward maps *C*. The SRs learn the predicted future occupancy of all features of the environment given the currently observed features (see methods) (**Figure 1B**). In contrast, the convolved reward maps *C* learn an average estimate of reward in each location using a filter that weights any rewards located either side of the current location (see methods) (**Figure 1C**). Importantly, at the same time as they are learning these maps, both algorithms also use Bayesian inference to compare the current observations from the environment with their current maps of the world to infer which context *z* they are most likely in. Specifically, FI compares the features *ϕ* observed by the agent prior to and following an action and compares how well the observed transition fits with the predictions encoded in each successor map *M*. In contrast, OI compares the observed convolved reward *cr* with the predicted *cr* from each convolved reward map *C*. Based on their current observations, the agents can either infer whether the current observations align well with an existing map; or if not, the agent can create a new map.

FI or OI cannot on their own be used to select actions, as FI does not learn about rewards and OI learns only a local representation of reward. As a result, in addition to this overall architecture, for each context a corresponding temporal difference *T D* map is learnt for action selection within each context (**Figure 1B,C**). For FI specifically, an alternative would be to learn a reward vector indicating which features are rewarded to allow for predicting the best current action. Here, we used *T D* maps for both strategies to (1) allow both FI and OI to use the same mechanisms for decision making to facilitate comparison of the two inference strategies and (2) isolate contextual inference from action selection. The *T D* maps learn value gradients across the locations and features of the environment, thereby allowing the agent to evaluate which action brings them closer to the reward (see methods for more detail). Once a context has been inferred, the context identity is passed to the corresponding context specific *T D* map to select the next action. Finally, the corresponding *T D* and *M* or *T D* and *C* maps are updated.

Importantly, the agents are not provided with any representations of the environment and do not receive any external context signals, aside from the environmental observations that indirectly indicate the hidden context. As a result, the main challenge for the agents is that they must simultaneously learn context-dependent representations, and use these representations to infer the current context and guide behaviour. As a simple comparative metric, if the algorithms learn accurate representations of both contexts, within the cued T-maze FI should be able to predict the current context based on the cue, whereas OI should be able to react to the absence of reward on the previously rewarded arm to infer the current context (**Figure 1D**).

### Feature and outcome inference outperform alternative strategies during cued switches in a T-maze

We first wanted to understand whether FI and OI can solve contextual inference tasks in principle, when contextual SNR is high. To do so, we tested whether they can solve the simple cued T-maze task, that has been readily shown to be solvable by other contextual inference algorithms [10, 19, 20]. Importantly, using the simple cued-T maze configuration described above provides a baseline, upon which we can then investigate how each model type is influenced by the amount of overlap between environments in different contexts by decreasing the SNR of the contextual signal. We will do so either by i) adding distractor cues around the predictive cue or ii) increasing the distance between the predictive cue and the choice.

In order to test whether agents and animals can perform predictive, cue-based inference in the cued T-maze we present trial types from each context in a random order, as in the computational and rodent literature [10, 19, 20, 46, 45, 47]. However, in laboratory settings an initial learning phase is required for animals to learn this task [46, 45, 47]. A common training technique to teach contextual inference tasks consists of using a blocked learning phase, where trials are arranged in blocks of the same trial type. This technique has been used in both the rodent and human literature in similar tasks [37, 48, 37, 49, 50], as well as in the computational literature, specifically for training Bayesian SR algorithms [14, 1]. Guided by this literature, we first trained FI and OI agents to perform the cued T-maze in a blocked learning phase, where trials were arranged as blocks of the same trial type, before moving agents on to the full task with randomised trials.

Once agents had completed 50 correct trials in one context, the context was switched such that the cue was different and the reward was located on the opposite arm. We trained agents on this version of the task for 10 blocks (500 correct trials). Next, we allowed these agents to continue to perform the task, and tested their performance on a further 10 blocks to see how quickly each algorithm reacted to changes in the trial type (**Figure 2A,B**). We found that both FI and OI models learnt to perform the block switches well, with OI requiring 2-3 attempts to adapt its behaviour following a trial type switch and FI requiring 1-2 attempts (**Figure 2C,D**).

**Figure 2.**
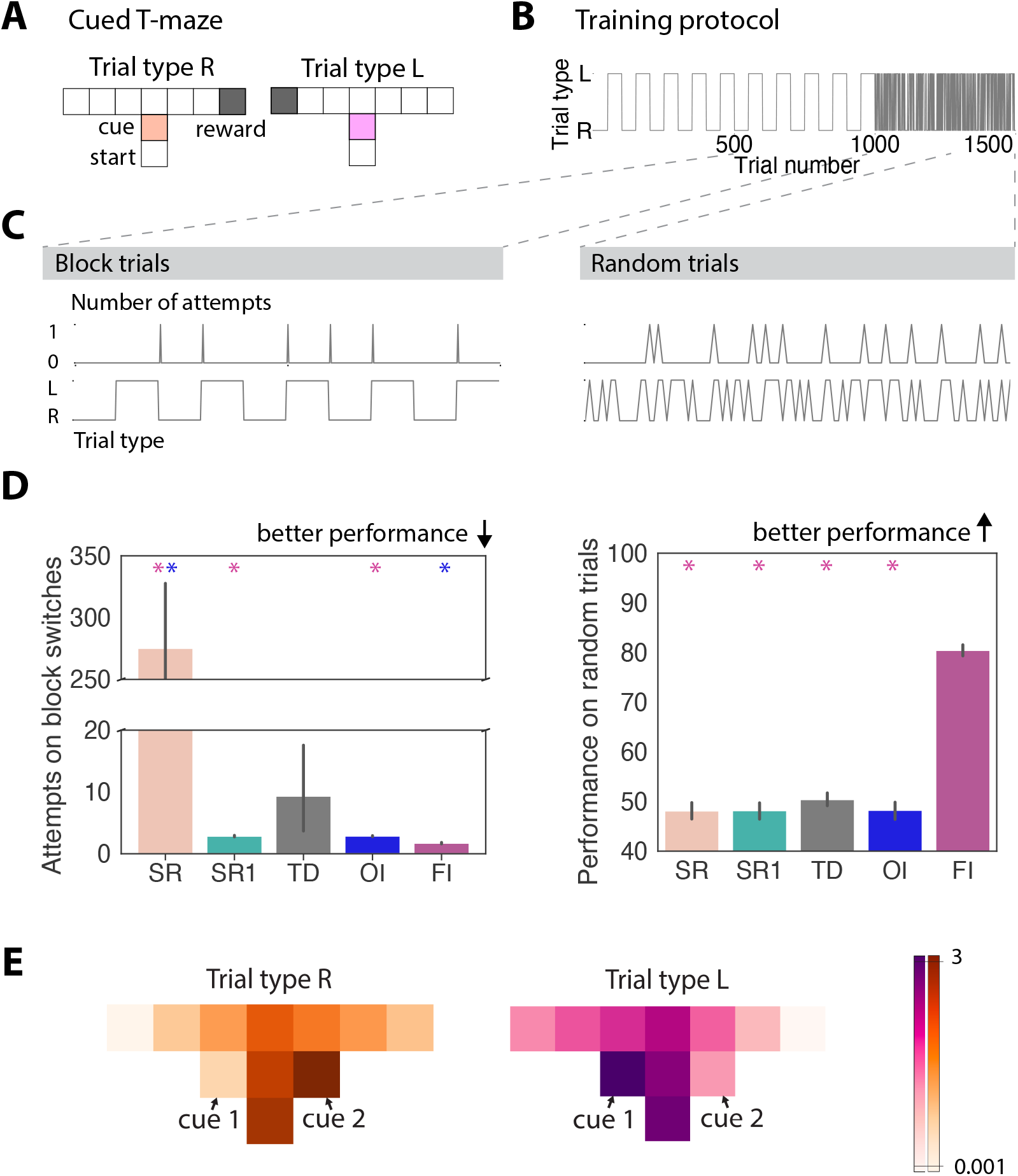
Feature and outcome inference learn distinct strategies to solve a cued T-maze. (A) Setup of the cued T-maze, showing distinct cues and reward locations for distinct trial types, (B) Training and test setup for the paradigm, showing trial identity for 1000 trials during block switches followed by 500 random trials, (C) Performance of an example feature inference agent on the last 10 block switches (left) and last 100 random trials (right) used to quantify their performance, (D) Performance of feature inference and outcome inference in comparison to other RL agents on block switches (left) and random trials (right), (E) Trial type specific SRs learnt by the feature inference algorithm showing distinct predicted future occupancy when the agent is in the starting state, averaged over all agents on the last 100 random trials (log-scale). Predicted future occupancy of locations is indicated within the T-maze and predicted future occupancy of cue 1 associated with trial type L is indicated to the left of the location it occurs in and cue 2 associated with trial type R is indicated to the right of the location it occurs in.

To confirm that Bayesian inference is necessary to perform contextual inference in the cued T-maze, we next compared the performance of these models with the performance of their individual components. We specifically compared the algorithms to an SR algorithm and TD algorithm that both learnt only one map of the environment. Within the SR algorithm, we compared two different versions i) where the reward vector was either updated incrementally (SR) or ii) where the learning rate was set to 1 such that rewards were learnt and unlearnt within one experience (SR1). We found that both feature and outcome inference performed better than the algorithms that do not use Bayesian inference on block switches (**Figure 2C,D**). This increase in performance of inference agents is due to the SR and TD algorithm both needing to gradually unlearn the reward location and relearn the new reward location following a trial switch. Even in SR1, following the absence of reward the agent must re-explore the whole environment to find the new reward location, as it has no memory of previous reward locations.

### Predictive inference is required to solve a cued T-maze

We next wanted to test the extent to which each algorithm could support predictive inference. To do this, we exposed each agent that had been trained on the block switch task described above, to a version of the task that required predictive cue-based inference within each trial, where the context (and hence appropriate behaviour) on each trial was chosen at random. Therefore in this instance, past outcome is not informative for behaviour, and so we reasoned that OI agents would not be ale to perform this version of the task. We let each agent perform an additional 400 trials before testing their performance over 100 further random trials (**Figure 2B**).

Consistent with this reasoning, FI agents could accurately predict the reward location based on the cue around 80 percent of the time, while OI agents could not. Indeed when we compared performance to all other model types (SR, SR1, TD), only FI algorithms were able to perform predictive inference (**Figure 2C,D**).

We next investigated what specific representation allowed FI agents to perform the task. We found that FI agents formed a distinct successor map for each context, including distinct representations of the cue identity and the associated behaviour that the agent follows. This can be seen by comparing the average predicted future occupancy of each feature when the agent is in the starting location across each context (**Figure 2E**).

Together these initial results confirm previous reports [42] that OI can learn to use an outcome-based strategy that is effective at solving tasks that contain blocks of repeating trials, but cannot solve random trial presentations, where the current best behaviour is independent of past outcomes. In contrast, FI can learn to use a predictive strategy, offering an improved performance on blocks of trials, as well as the ability to solve random trials.

### Performance of feature inference degrades as contexts become more similar

Our results so far suggest that in principle, FI is well-suited to perform predictive inference in the cued T-maze task. We next looked at the effect of decreasing the contextual SNR, by increasing environmental overlap between contexts. We tested this with two complementary manipulations to the task structure. First, we created a series of different mazes, where we incrementally added an increasing number of ‘distractor’ features that are present in both contexts around the cue (**Figure 3A**) [51, 52, 53]. In other words, while in the base task the cues may have been A vs B, a version with 3 distractor cues has 4 cues in each maze in total, three of which overlap: XYZA vs XYZB. To compliment this, we also created a series of mazes where instead we incrementally increased the distance between the cue and the choice-point, without changing the number or identity of the cues (**Figure 3D**) [44, 46, 45]. In this task the increased overlap between the two contexts arises from the increasing number of locations in the centre arm, that due to their lack of distinguishing features are highly similar across contexts.

**Figure 3.**
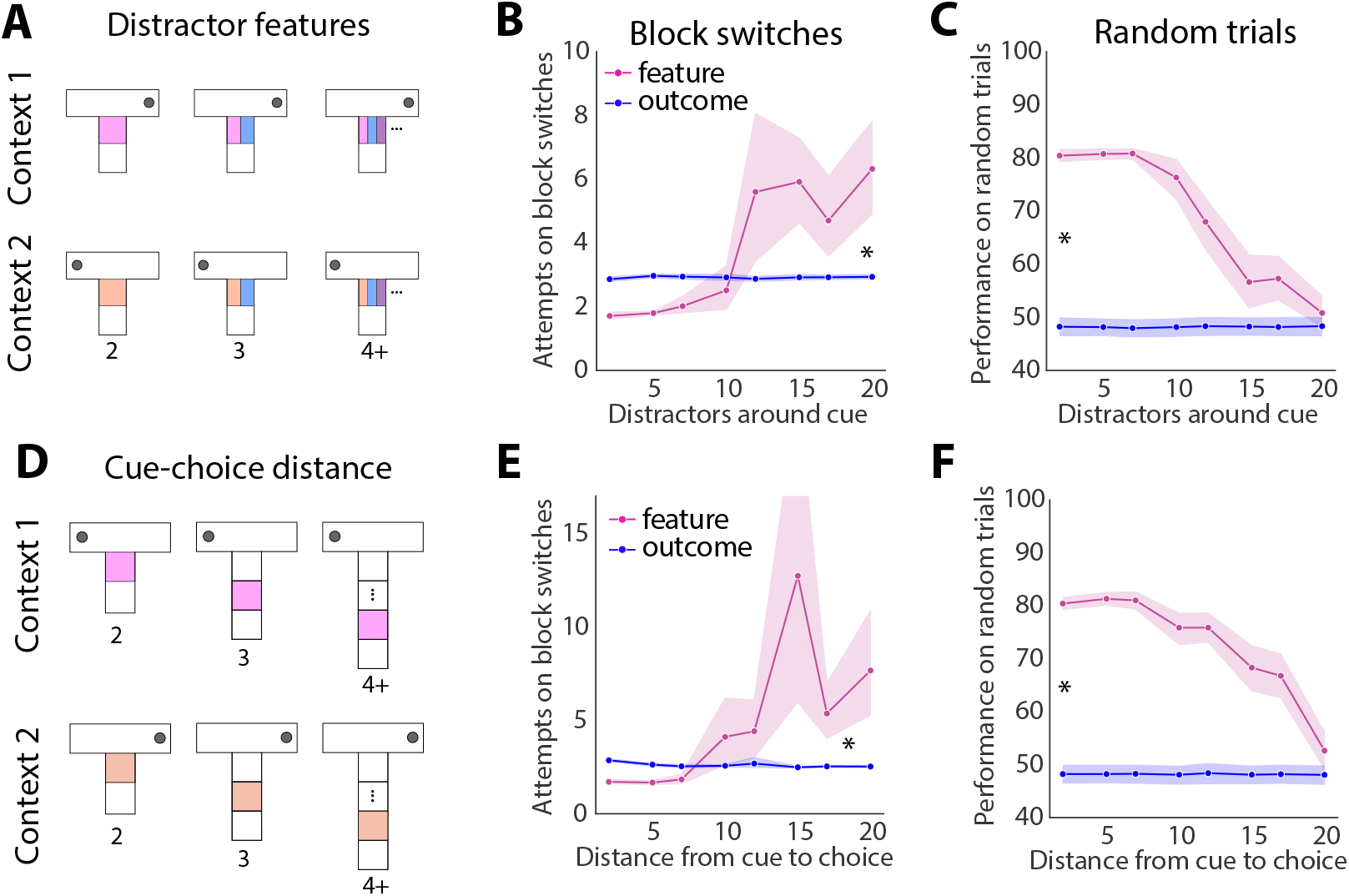
Increasing overlap between contexts reduces performance of feature inference, but not outcome inference. (A) Decreasing the contextual SNR by adding multiple overlapping features around the cue in each context or (D) extending the distance between the cue and the choice point, (B, E) Performance of feature and outcome inference as contextual overlap increases on block switches and (C, F) random trials

Using this approach, we found distinct effects of distractors both across stages of the task, and across FI and OI agents. These differences were independent of the means by which the overlap increased. During the block switching stage of the task, the number of attempts required by FI agents to reverse increased consistently with increasing environmental overlap between contexts (**Figure 3B,E**). In contrast, we found that the number of attempts to reverse on block trials using OI was extremely consistent independent of the amount of environmental overlap between contexts (**Figure 3B,E**).

Similar to the block switching stage, the performance of FI on random trials decreased steadily to chance with increasing contextual overlap (**Figure 3C,F**). This was in contrast to OI agents remaining unable to perform above chance for any of the distractor combinations. Together, these results indicate that during learning, OI is robust to decreases in contextual SNR, resulting from increasing overlapping features. This suggests that OI may be well-suited to provide a context signal to FI agents that otherwise struggle during learning with low SNR.

### Supporting feature inference with outcome inference during learning rescues performance at low SNR

Our data suggest that FI agents increasingly struggled to perform the T-maze task as the two contexts became more similar. In contrast, OI agents were able to stably learn to separate the two contexts, independent of their overlap. Therefore we reasoned that supporting FI with OI during learning may overcome the limitations of FI, by supporting more accurate initial learning.

To test this, we built a joint inference architecture where OI supports FI during learning (**Figure 4A**). To implement this, we began by generating joint agents containing both FI and OI algorithms, and allowing the algorithms to interact with one another only during the initial learning phase - for the first 200 trials of the task. During these trials the agent created a ‘joint estimate’ of which context is most likely by calculating the average of the probability according to the FI and the OI algorithms (**Figure 4B**). We chose the average over the first 200 trials as a proof of principle experiment, as it allows us to investigate whether the simple metric of averaging implemented for very few trials at the start of learning is sufficient for improving performance, and leaves room for more complex methods in the future. This jointly inferred context identity is then used to select the optimal choice and to update the corresponding outcome and feature inference maps. After the first 200 trials, the agents then revert to a solely FI strategy, and no longer incorporate information from OI.

**Figure 4.**
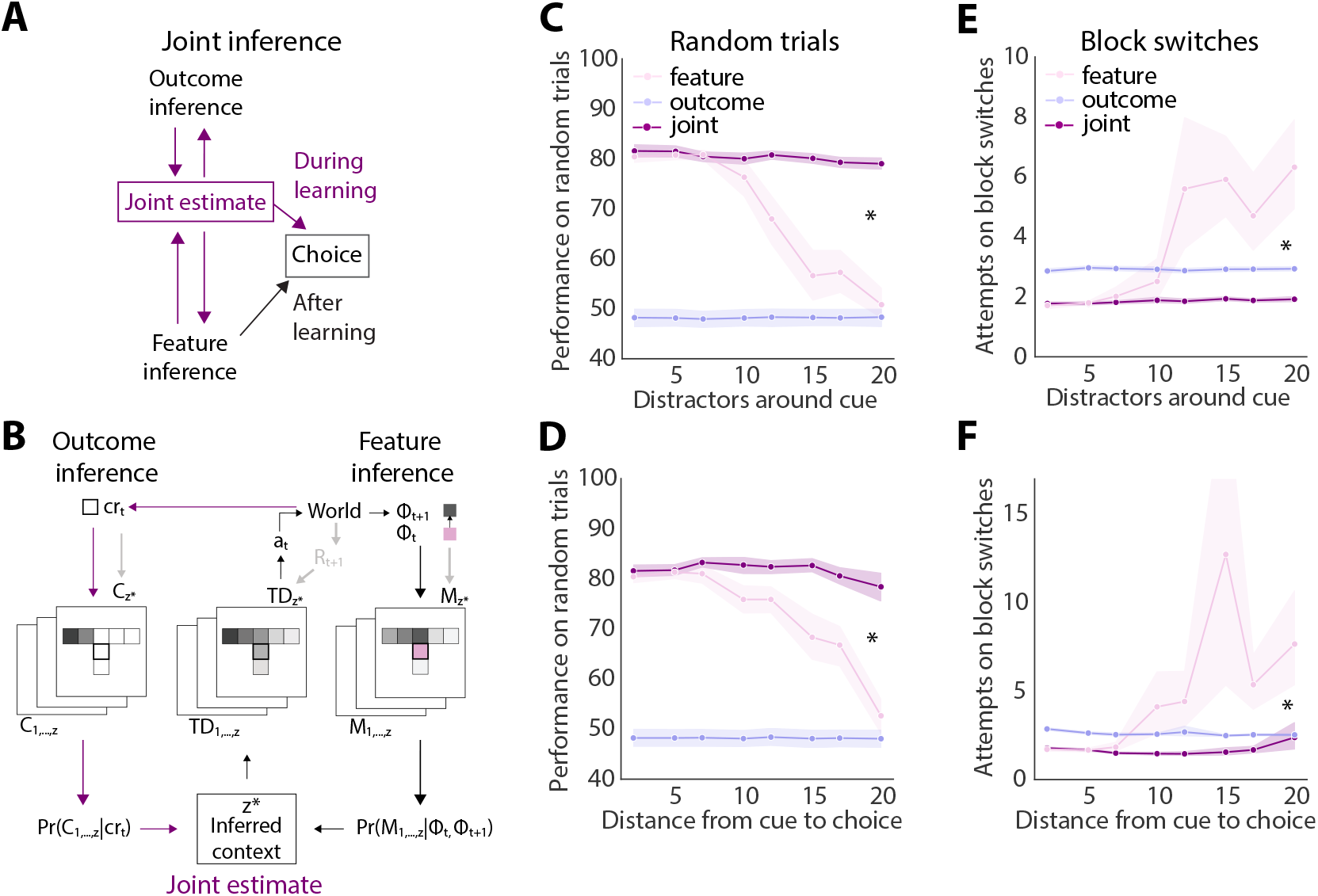
Supporting feature inference with outcome inference during learning rescues performance with increasing contextual overlap. (A) Schematic of joint algorithm showing that during learning outcome inference is used to generate a joint context estimate, (B) Algorithmic implementation showing how the joint estimate is determined during learning, (C) Performance of joint inference on random trials as contextual overlap increases via distractor features around the cue or (D) increasing cue-choice distance, (E) Performance of joint inference on block switches as contextual overlap increases via distractor features around the cue or (F) increasing cue-choice distance. **Figure 4—figure supplement 1**. Inconsistent performance of algorithms using other ways of combining outcome inference and feature inference during learning.

When we tested this joint inference model, in contrast to FI alone, we found this algorithm had stable performance across all contextual SNRs, both in the case where SNR decreased with increases in distractor features and in the case where SNR decreased with increases in distance between the cue and the choice (**Figure 4C-F**. Crucially, although we only allowed the interaction of the two agents for from first 200 trials of the block switches, this stable performance lasted throughout the remaining block switches, and also resulted in stable performance during random trials. Thus, supporting FI during learning with OI rescues deficits in performance with low SNR, even after input from the OI algorithm is no longer present.

Importantly, the ability to incorporate OI into FI estimates can occur in multiple ways - not just via real-time access to both estimates to drive contextual inference. Therefore, we also tested whether this ability to support feature inference could be achieved using multiple alternative implementations of the interaction. In particular we focused on a model that uses replay of contexts inferred by outcome inference after each trial (**Figure 4-S1A**), and a model that uses ideal observer estimates of the current context (**Figure 4-S1B**). In both cases we found that such approaches did not consistently overcome the deficits of feature-based learning. Therefore, while multiple mechanisms could support this joint estimate, it seems that real time access to both outcome and feature based representations is particularly beneficial for accurate contextual inference at low SNR.

### Access to outcome inference stabilises the learning of feature-based representations

Our data confirm that the presence of OI information during learning greatly facilitates the use of FI, even long after this OI information is no longer available. We next wanted to investigate *how* the addition of OI to the FI algorithm supported this facilitation of learning. We found that in the block learning phase, FI, but not OI or joint inference, struggled to maintain performance as environmental overlap between contexts increased. We reasoned that the distinction between algorithms is that in FI it is necessary to learn to predict the future occupancy of all of the features in the environment, whereas OI only learns to predict one feature - the reward. As overlap between contexts increases, the features of the environment become more similar between contexts, but the observed rewards do not. Therefore, when contexts are highly overlapping, early in learning FI agents will be likely to assign observations to the incorrect context, thereby reducing their ability to distinguish between the different contexts. As a result, specifically the first contextual switch could be impacted by increasing contextual overlap, which could influence the long term representations of the FI model, by degrading the difference between the representations.

To test this, we investigated the first time the agent experienced a switch in trial type (the first switch from context A to context B in the block switch stage). We reasoned that even on this first switch, when overlap between contexts is high, the FI algorithm would incorrectly incorporate new observations in the original context, rather than create a new context from these new observations. Consistent with this, we found that agents using FI displayed a large increase in the amount of incorrect updates to the original context representation as overlap increased. In contrast, and consistent with our previous data, this incorrect updating did not occur when using OI or joint inference, and these differences between FI, OI and joint inference were independent of whether overlap was increased by increasing the number of distractor features, or increasing the distance form the cue to the choice point (**Figure 5A**).

**Figure 5.**
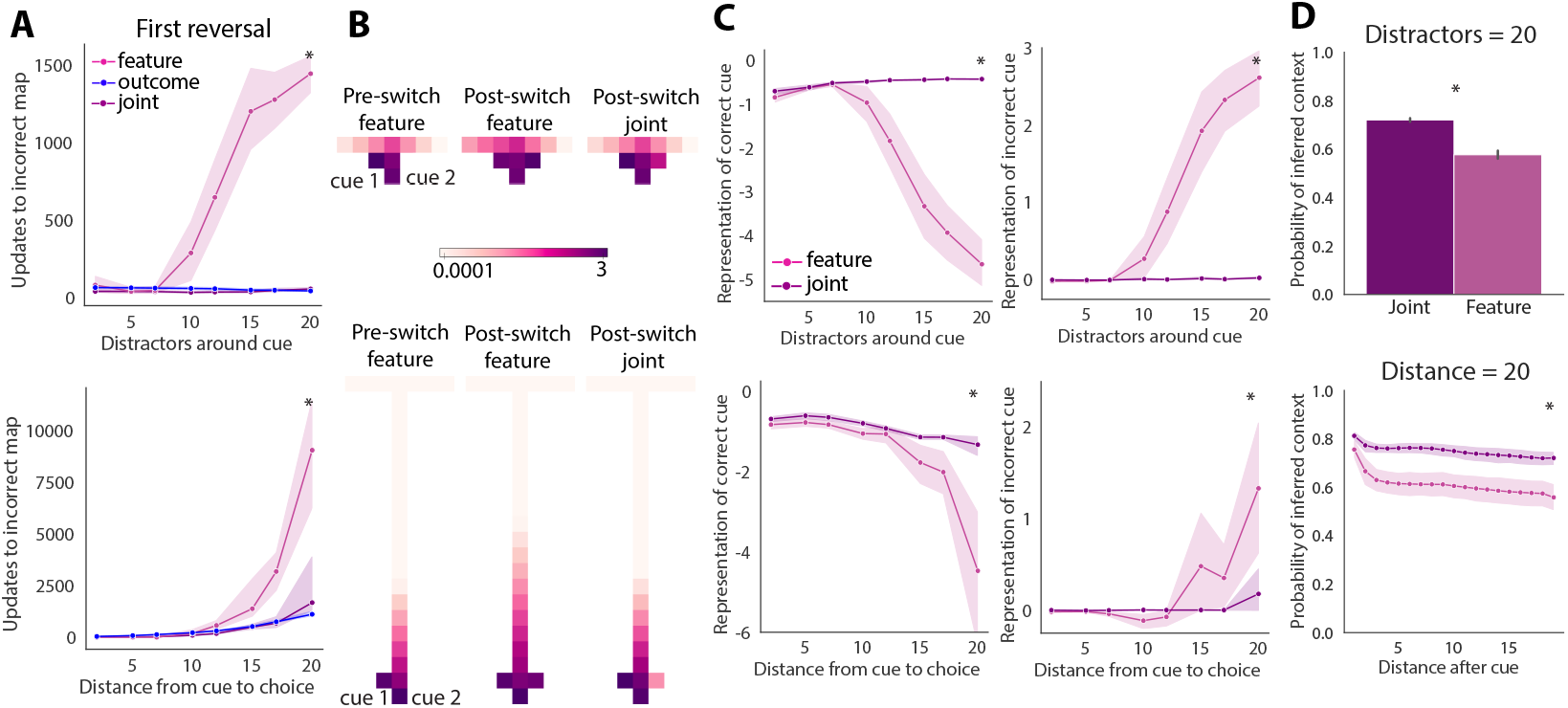
Supporting learning with outcome inference improves initial formation of context-dependent representations leading to long-lasting improvement in performance. (A) Number of incorrect updates of the context map during the first trial type switch, (B) Evolution of the average context map’s predicted future occupancy when the agent is in the starting state (log scale), at the end of the first block and the end of the second block of trials for joint and feature inference agents, (C) Underrepresentation of the correct cue (left) and overrepresentation of the incorrect cue (right) on the average context maps used on random trials. (e) Confidence in inferred context identity following the cue on random trials. **Figure 5—figure supplement 1**. Inconsistent performance of algorithms using methods other than outcome-inference to support learning.

We next investigated what influence this incorrect updating had on the representations of context. To do this, we plotted the predicted future occupancy of each environmental feature when the agents are at the start of the maze both before and after the first trial type switch (**Figure 5B**). We found that at the end of the first block, preceding the first trial type switch, there was a clear distinction between the expectation of cue 1 and cue 2, despite multiple distractors (**Figure 5B**). We then compared the effect of the first trial type switch on the representation of both cues in the FI and joint inference algorithms. Interestingly, when using FI, the incorrect cue became incorporated in the original context representation (note high expected cue 2, **Figure 5B**). However, this was not the case when FI was combined with OI in the joint algorithm (note that cue 1 continues to be the most expected, **Figure 5B**). Therefore consistent with our prediction, FI agents incorporate new observations into the wrong representation after the first switch in context.

Interestingly, we found that this degradation of context-specific maps when using FI was long-lasting: on random trials the FI agents had a decreased representation of the correct cue and an increased representation of the incorrect cue in both context representations as the amount of environmental overlap increased. This was despite these trials occurring many hundreds of trials after the first trial type switch. This long lasting degradation was also mitigated by use of a joint agent, despite again the access of this agent to OI information being removed many hundreds of trials earlier (**Figure 5C**).

This degradation of context-specific maps in FI agents had a marked behavioural effect - it resulted in a decrease in the inferred probability of the context following the cue on correct trials, indicating a larger chance of inferring the incorrect context (**Figure 5D**). Therefore, we found that increasing environmental overlap between contexts impairs the ability of FI to learn distinct context representations. This deficit can be compensated for by integrating outcome inference during learning.

Importantly, this specific implementation of the FI model makes a number of mechanistic assumptions, and so we also tested multiple alternate methods for making inference of a new map easier for FI when there is a large amount of overlap between context features. First, we gave the FI agent access to an additional map filled with a random-exploration SR (learnt through random exploration of the environment - see methods) when each new context was first introduced [54]. This removes the requirement to create a new, unexplored contextual map upon inference of a contextual switch, which we hypothesised could in some cases decrease the likelihood of switching representations. However, this approach did not improve performance of the FI agent, as the lack of directionality in the random-exploration SR led to a poorer fit with the new experiences than the previous context map (**Figure 5-S1A**). Second, we tested the influence of adding the reward as a feature, which could intuitively be thought to further distinguish the two contexts based on reward location [14]. Interestingly, this approach further increased the amount of environmental overlap between different contexts during learning and thereby unreliably impacted performance (**Figure 5-S1B**). Lastly, we found that blocking access to the incorrect arm during training, forcing the choice of the agent towards the rewarded arm, a strategy often employed during rodent experiments [44, 46, 45], also did not consistently improve performance of the feature inference algorithm (**Figure 5-S1C**). To summarise, the most effective means we found to support the formation of distinct context maps in FI is by integrating outcome inference during learning.

Together, our results indicate that as overlap increases between contexts, the ability of FI agents to perform predictive inference decreases. This is similar to previous observations of decreasing performance of RNNs as overlap between contexts increases in similar tasks [10, 19, 20]. Interestingly, we found that the deficits in FI could be rescued by using OI to guide initial learning of feature representations. Specifically, we found that OI could be used to stably identify changes in context during learning, allowing for the FI algorithm to learn distinct representations of each context, resulting in long-lasting improvements in behaviour.

### Joint agents produce internal variables similar to hippocampal splitter cells

A core goal of our study was to frame the interaction between these algorithms in a way that was interpretable from a neuroscience perspective. Neural activity in the CA1 area of the hippocampus (HPC) has been proposed to exhibit an implementation of SRs, where the activity of individual neurons represents the predicted future occupancy of a specific location [35, 55, 56]. Multiple neuroscience studies have shown that a signature of HPC activity in context-dependent tasks is the presence of so called ‘splitter cells’: cells that fire differently in the same location in an environment depending on the inferred context [57, 58, 59, 60]. These cells are thought to allow for similar locations in different contexts to be represented with distinct neural activity [25, 37, 1]. We hypothesised that agents utilising FI (therefore both FI and joint inference agents) would produce activity reminiscent of splitter cells, as in these models different SRs are learnt and used to guide behaviour in each of the two contexts of the task [42, 1]. To investigate this directly, we estimated simulated ‘cell firing’ in each of our agents, using the future predicted occupancy of a chosen location, weighed by the probability of the context map being inferred. We then calculated this firing for each context map at every location in the environment to create a set of ‘place cells’ [35] for each agent.

We then used this metric to compare simulated cell firing along the shared central arm of the maze - similar to classic ‘splitter cell’ papers [57, 58, 59]. For each simulated cell with a firing field in the central arm, we found the location and context in which its ‘activity’ was highest, and compared this ‘activity’ to the same location in the other context. When we did this, we found that both joint and FI models produced patterns highly reminiscent of ‘splitter cells’ (**Figure 6A**,**B**). Therefore, similar to previous work [42, 1, 34, 11] in our task, FI models can reproduce classic HPC splitter cell experiments.

**Figure 6.**
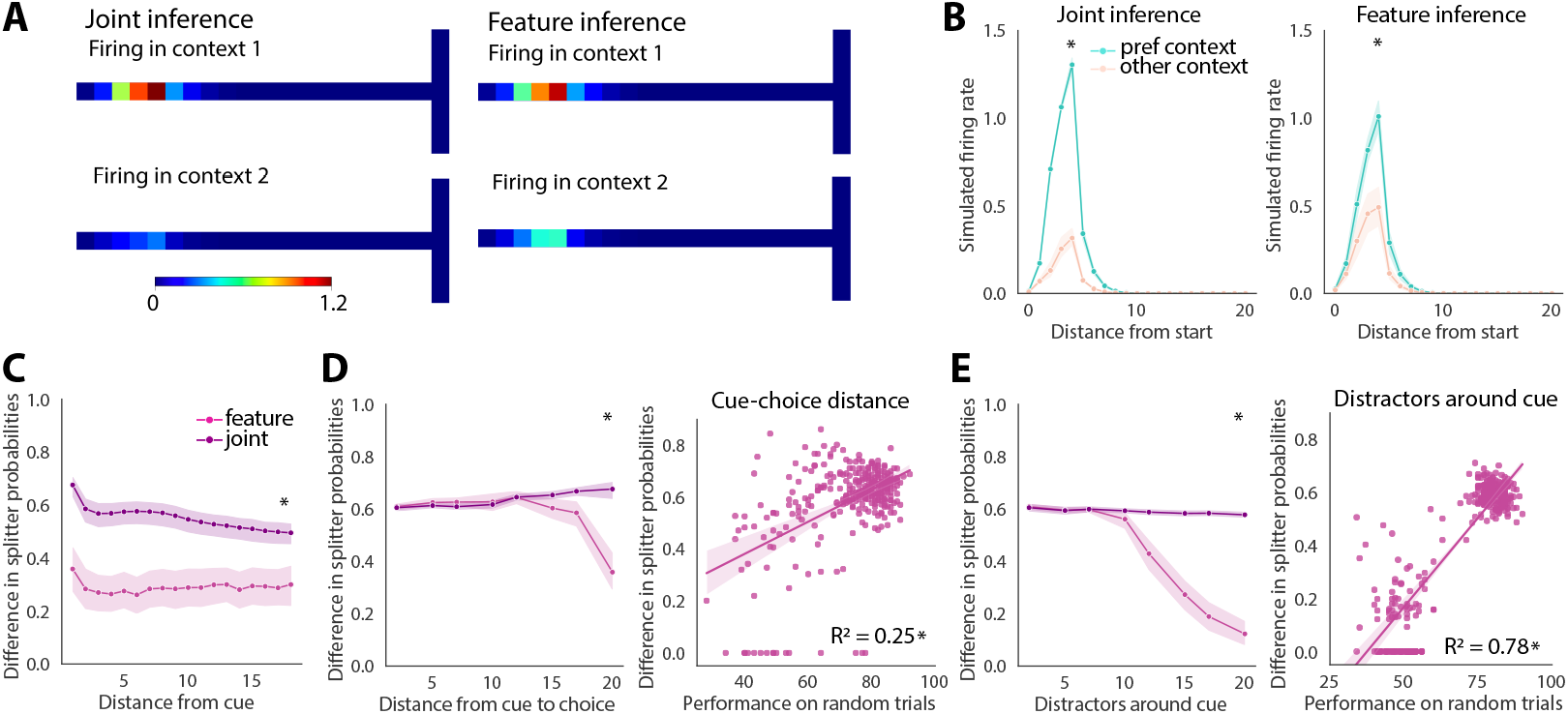
Removing support from outcome inference during learning mimics experimentally observed splitter cell loss. (A) Simulated cell firing of a cell representing future predicted occupancy of states along the overlapping central arm on correct random trials across all agents for cue-choice distance 20, (B) Quantification of simulated cell firing along the central arm in its preferred and non-preferred contexts, (C) Evolution of the difference in splitter probabilities along the central arm, (D) Impact of increasing cue-choice distance on the difference in splitter probabilities (left) and correlation between performance on random trials and the difference in splitter probabilities (right). (E) Same as (D) but for distractor features.

**Figure 7.**
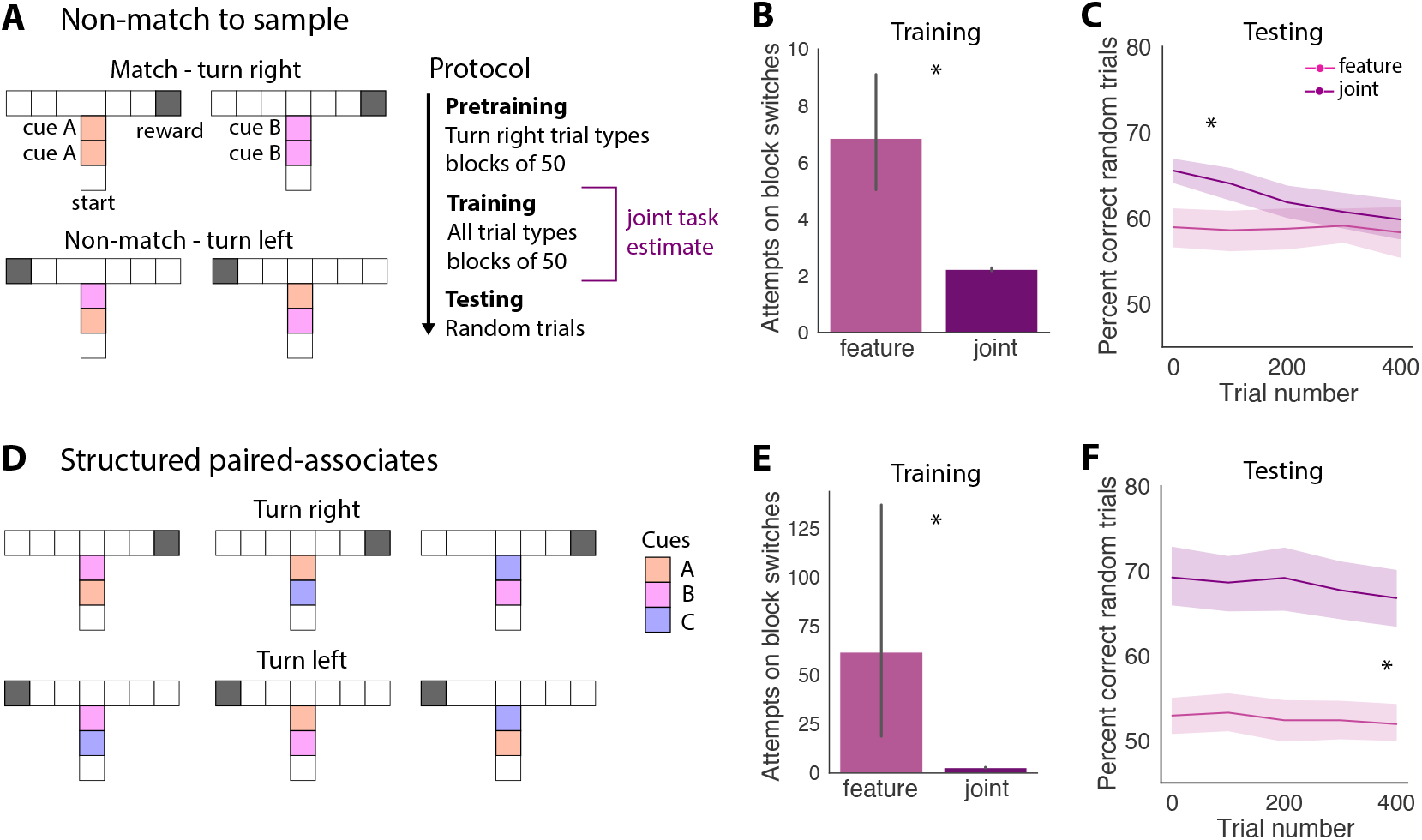
Supporting feature inference with outcome inference during learning improves performance on cue-discrimination tasks. (A) Schematic of the non-match to sample task, where trials with the same cue repeated twice require a different response than trials where two differing cues are presented (left) and training protocol showing that outcome inference is used to support feature inference during the training phase (right), (B) Performance of feature inference and joint inference on the last 10 block switches, (C) Performance of feature inference and joint inference on random trials, (D) Schematic of the biconditional discrimination task, where the meaning of the second cue depends on the identity of the first cue, (E) Performance of feature inference and joint inference on the last 10 block switches, (F) Performance of feature inference and joint inference on random trials.

### Removal of outcome inference mimics loss of HPC splitter cells after PFC lesions

We next wanted to investigate whether the interaction between FI and OI may behave in a similar way to the interaction between PFC and HPC. Consistent with this idea, PFC is often thought of as performing outcome inference [6, 39, 37] and its input to HPC is crucial for accurate learning [25, 26, 27, 28]. Moreover, if PFC input to the HPC is eliminated, there is a dramatic loss of splitter cells in the HPC [25, 37], suggesting that loss of PFC input degrades the representation of context in HPC. In our framework, during learning the joint inference algorithm could be described as using contextual estimates based on OI calculated in PFC, and integrating these with contextual estimates based on feature inference in HPC. We therefore wanted to investigate if removing OI from the joint algorithm during learning could replicate the loss of splitter cells in HPC caused by removal of PFC input.

To investigate this, we returned to our simulated splitter cell analysis described in **Figure 6A,B**. We compared the properties of simulated cell activity from the FI algorithm with or without the addition of OI. As before, we found that in the full joint inference algorithm cells showed a large difference in firing between their preferred and non-preferred contexts (**Figure 6A,B**). In contrast, in the algorithm without outcome inference, this difference in firing was greatly reduced (**Figure 6A,B**).

We next quantified this loss of splitter cell-like activity more generally by looking at the difference in the probability of firing of each cell in its preferred versus non-preferred context. Using this metric, we found large differences in activity in each context along the entire stem of the maze in agents utilising joint inference. Consistent with our previous findings, we also found that these differences were consistently lower in agents without access to OI (**Figure 6C**). Further, we found that while increasing environmental overlap between contexts had minimal effect on splitter-like activity in joint inference agents, increased overlap led to a large reduction in this difference in activity in agents without OI (**Figure 6D,E**). Again, this degradation was similar irrespective of whether overlap was increased using distractors or distance between the cue and the choice point. Consistent with this, we found that agent-by-agent, decreased behavioural performance without OI was strongly correlated with decreased differences in splitter-like activity (**Figure 6D,E**).

These experiments suggest that the joint inference model provides a framework describing HPC activity during behaviour requiring use of hidden contexts, and also the contribution of PFC to the learning of this behaviour. Using this framework, we found evidence to suggest that the strength of splitter cell specificity indicates how different context representations are from one another. In turn, this also indicates how well the agent will perform on tasks involving predictive inference.

### Outcome inference supports learning of a delayed non-match to sample task

In our results so far, we have shown that supporting FI with OI during learning generates a robust and long lasting basis for performing predictive inference. Until now however, our experiments have been limited to a relatively simple T-maze task. Although this task is widely used in the neuroscience literature, we next wanted to investigate if the interaction of FI and OI may be a more general mechanism to facilitate performance on contextual inference tasks. To do this, we focused on two alternate tasks - first a classic delayed non match to sample task, which has been used for decades to study PFC and HPC function [61, 62, 63, 64], and second a more complex structural learning task, that has been proposed to isolate a core function of the HPC in associative learning [65, 66, 67, 68].

First, we investigated a delayed non-match to sample task - a behavioural task strongly dependent upon activity in both PFC [26, 69, 70, 71, 64] and HPC [72, 61, 73, 62, 63]. In this task, two cues are presented one after the other in time. If the cues do not match (i.e. the first and second cue differ from one another) the agent must make one choice, for example, ‘go’, or choose right, whereas if the cues match the agent must make a different choice, for example, ‘no-go’ or choose left (**Figure 7A**) [69, 70]. Therefore, we implemented this task using a similar layout as our T-maze, where two cues were presented one after the other as an agent traversed a central arm. Subsequently, the agent had to make a choice to turn left or right, based on the preceding cues: matching cues (A-A or B-B) cued a right choice, while non matching cues (A-B, or B-A) signalled a left choice. Based on the rodent literature, learning requires that animals are pretrained first on trials associated with only one outcome, before adding in trials associated with the other outcome [26]. To be able to directly compare with this literature, we therefore pretrained agents first on trials where the reward was located on the right arm using blocks of trials of each trial type (i.e. A-A or B-B) (6 blocks each, consisting of 50 trials), followed by training the agent on blocks of all trial types (A-A, B-B, A-B or B-A) (5 blocks each, consisting of 50 trials). Finally, we tested whether agents learnt to predict the reward location by testing agents on 500 randomly selected trials.

Based on our previous results, our prediction was that the addition of OI to FI during initial learning would facilitate the performance of FI during and following learning. Therefore, we built a joint algorithm where OI supported FI during the block training phase. We first compared the performance during the block training phase of the joint inference algorithm, where OI supported FI, to the FI algorithm without OI support. Similar to our previous results in the T-maze, we observed that during the delayed non-match to sample task the use of joint inference dramatically improved performance during training (**Figure 7B**). This improvement also led to improved performance when tested on random trials. Interestingly however, in contrast to the cued T-maze, the effect was not long lasting, and only apparent for the first 200 trials (**Figure 7C**). These data suggest that supporting FI with OI during learning also improves the formation of distinct context-representations for context-dependent decision making during a delayed non-match to sample task. Additionally, the overlap between tasks leading to degradation of performance over time when OI was no longer present, indicates that additional support may be necessary to ensure stability of representations after learning. Notably, this result is highly consistent with data observed in recent animal work, where PFC activity is required for accurate learning of such tasks [26], and further suggests a role of PFC activity may be to provide stable contextual estimates based on outcome inference to HPC as feature representations are being constructed.

### Outcome inference supports learning of a structured paired-associates task

While delayed non-match to sample tasks have been widely historically used, a number of alternate ways to solve such tasks, have resulted in it falling out of favour as a means to study contextual behaviour. Recently developed structured paired-associate discriminations have been designed with the aim to mitigate such issues [67, 68]. Much like a delayed non-match to sample task, these tasks consist of trials each with two cues separated in time. However, to prevent the use of alternative strategies, each trial type is formed from a pool of 3 cues (A,B and C) that are combined in a specific way. Firstly, they are ordered such that the meaning of the second cue is dependent on the first (e.g. A-B is different from C-B). Secondly, the identity of the first cue is also uninformative without the second cue (A-B is different from A-C). Finally, the order the cues are experienced in is also critical (A-B is different from B-A). These trial types are arranged into 6 combinations such that the agent must use a combination of the two features in order to solve the task (**Figure 7D**).

We next investigated if FI agents could solve this complex task, and whether this performance was facilitated by the addition of OI during learning. To test this we trained agents using the same stepwise approach as the non-match to sample task, where we first pretrained in blocks of only ‘turn right’ cue combinations (3 blocks each, consisting of 50 trials), followed by blocks of both ‘turn right’ and ‘turn left’ cue combinations (3 blocks each, consisting of 50 trials) (**Figure 7D**). In this complex task, we found that agents with FI alone struggled to perform well - these agents struggled to solve the block stage and plateaued at only slightly above chance performance when presented random trials. However, we again found that supporting FI with OI during training dramatically improved performance. This was apparent both during training (**Figure 7E**), and during random trials (**Figure 7F**). Interestingly, contrasting with the non-match to sample task, but consistent with the T-maze task, the support provided by OI during learning was essential and led to long lasting improvements on random trials, where agents lacking the support of OI performed poorly, with performance remaining near chance. Together, these results indicate that during learning OI can support the formation of stable FI representations across a range of neuroscience tasks.

## DISCUSSION

Most of the decisions that we make on an everyday basis are context-dependent decisions. However, we still do not understand how context-dependent learning is supported by the brain, nor do we have algorithms that can perform context-dependent learning without a constant, externally provided context signal. To address this issue, we first investigated how well current predictive algorithms can learn to perform contextdependent decision tasks. We found that as contexts become harder to distinguish from one another, predictive algorithms in general struggle to learn distinct contextual representations and thereby struggle to achieve good task performance. Focusing on a Bayesian model-based predictive algorithm, we found that supporting this predictive algorithm with outcome inference during learning stabilises learning across a wide range of contexts with increasing similarity, and across a range of distinct behavioural tasks used in neuroscience studies. Finally, we found that these algorithms reproduce observations from neuroscience experiments. Specifically, our results suggest that OI supporting FI during learning may be viewed as representing a model where context signals inferred using outcome inference are provided by PFC to HPC during learning, to support the formation of accurate predictive feature representations. Our results indicate that this framework replicates the behavioural- and cellular-level results of circuit-manipulations in animal studies across multiple behavioural paradigms, where inhibiting PFC input to HPC during learning prevents learning and produces signature loss of context-specific ‘splitter cell’ activity. Our results indicate that predictive approaches when supported with outcome inference during learning, make up a resilient system for predicting the optimal behaviour in context-dependent decision tasks. Together, our results offer a framework for interpreting behavioural strategies and cellular activity from neuroscience experiments during learning of context-dependent behavioural tasks. Our future work will aim to use this framework to investigate how the brain supports context-dependent learning.

### Implications for understanding how the brain performs contextual inference

A key caveat of a number of the most promising models of contextual inference, is that such models require an explicit context signal to be provided in order to successfully to support learning [23, 22, 21, 24]. Here we propose how such a context-signal might be simultaneously learnt during experience, removing the need for this provision. Moreover, we show that this approach can learn to solve multiple behavioural tasks commonly used in animals in neuroscience studies. In the future, using these models may allow us to tease apart the extent to which different circuits or brain regions carry distinct functions to support context-dependent learning. For example, our model predicts that for learning about abstract concepts, PFC may provide outcome-based support to HPC during learning [27, 25, 26, 28]. On a more conceptual level, our models provide a framework that can begin to help us understand how the interplay of different brain regions that each focus on different details of our experiences, can together guide learning.

### Limitations of model architecture

It is important to point out that because our algorithms are designed based on first principles, we make a number of assumptions about how the agents explore and learn about the environment.

Firstly, we assumed that the agents learn using a random walk, however a more targeted strategy, for example Lévy flight, could make learning of representations easier [74]. Secondly, we focused on *ϵ*-greedy explore-exploit strategies in this study, and it is important to note that alternative strategies could also aid in improving learning speed [42, 75]. This is because the way that the agents learn about the environment is extremely important to how quickly they learn and how well they perform on the paradigm. Even the differences in the random-walk each agent uses to explore here lead to large differences in how well the task is learnt, visible in the variance around average performance. As a result, improving the speed of learning individual context representations may reduce interference between context maps.

Additionally, the way in which the outcome and feature inference agents identify different contexts depends on the initialization of parameters for the algorithms. As a result, different behavioural tasks will require the use of distinct parameters. For outcome inference, the filter length determines how distinct different reward locations need to be to be recognised as belonging to distinct contexts. For both algorithms, the optimal prior will depend on the number of distinct contexts. Despite this, it is worth noting that these parameters are all interpretable from a computational and neuroscience perspective, and remain far fewer than those needed to fit an equivalent neural network approach.

Furthermore, the current design of our algorithms require the trials during learning to be presented in blocks, where multiple trials are presented in a row from each context. If tasks did not have this structure the facilitation by outcome inference would not be possible within our current joint inference framework. We chose this design as it is common in neuroscience studies, where often both animals and humans have been shown to learn best when initially presented with different contexts in blocks of trials. As a result this design is highly applicable for a wide range of tasks currently used within neuroscience [48, 37, 49, 50]. Additionally, alternative methods that do not require this structure will be interesting to examine in the future. For example, replay of experience at the end of each trial has been suggested to be able to support learning without such block structure - by essentially allowing for a compressed repeat of the experience to facilitate learning [42, 76, 77, 78, 79]. Although our implementation of replay here did not provide consistent improvements in learning within the block structure used here, it remains to be seen whether it could be used in situations where such a block structure is not present.

Finally, in the current implementation of our model, we manually determine when outcome inference should be used to support feature inference - by manually defining this interaction for the first 200 trials. In the future, it would be interesting to see how this can be automated, for example, by weighing the current utility of outcome versus feature inference. One way to achieve this could be to use the discrepancies in predictions between each model, as recently proposed for combining model-based and temporal difference learning to solve contextual inference problems [43]. This addition will also allow for investigation of how the brain may determine when it is necessary to provide support to predictive approaches using outcome-based information.

## Conclusion

Overall, our results indicate that models that support feature-based inference with outcome-based inference during learning provide a powerful framework for understanding how we learn to make decisions to support contextual inference.

## METHODS AND MATERIALS

### Behavioural task design

#### Cued T-maze

We modelled the T-maze as consisting of discrete spatial locations in the form of one-hot vectors, on top of which we added cues as distinct features. Each arm of the maze is made up of three distinct locations resulting in a state-space *S* ∈ [*s*_1_, *s*_2_… *s*_9_]. Observations emitted at the start of the stem of the maze then take the form of *ϕ*(*s*_1_) = [1, 0, 0, 0, 0, 0, 0, 0, 0, 0, 0]. The predictive cues are overrepresented 4x in comparison to the locations and are present at the second location on the stem resulting in the observation *ϕ*(*s*_2_) = [0, 1, 0, 0…, 4, 0] when in one context/trial type and *ϕ*(*s*_2_) = [0, 1, 0, 0, …, 4] when in the other context/trial type. The agent moves between states by selecting an action from the set of actions available to them *A* ∈ [*up, down, lef t, right*]. If the selected action involves running into a wall, the agent remains in its previous state. When the agent reaches the end of the correct arm, it receives a reward.

#### Non-match to sample task and structural discrimination task

For the non-match to sample task and structural discrimination task we used a modified version of the T-maze design. Behaviourally these tasks are usually implemented as head-fixed tasks, where animals either must choose to lick from distinct left or right spouts [69, 70] or preemptive licking on a set of rewarded trials and no preemptive licking on a set of non-rewarded trials is used as the read out for whether animals have understood the task [26, 80, 81, 82]. Within our model the second case can be interpreted as rewarded trial types being equivalent to receiving reward at the end of the right arm of the maze and non-rewarded trial types being equivalent to receiving reward at the end of the left arm of the maze. As a result actions taken that move the agent to the right can be thought of as preemptive licking, whereas actions taken that move the agent to the left can be thought of as not licking preemptively. Additionally, we only allowed the action *A* ∈ [*up*] on the central stem of the maze to mimic the agent being presented with cues rather than physically moving between them. For the non-match to sample task, the predictive cues are overrepresented 2x and for the structural discrimination the predictive cues are overrepresented 4x.

#### Training of algorithms

For the cued T-maze task, we trained agents on 20 block switches consisting of 50 trials each, such that the agents see each trial type 10 times. To test predictive inference, we then trained the agents on 500 random trials. For the joint inference algorithm, we used the joint prior and joint task estimate for the first 200 trials, which is equivalent to the first 4 block switches.

For the non-match to sample task, we pretrained agents alternating between blocks of the rewarded trial types until each trial type had been seen 6 times, with each block consisting of 50 trials. We then trained the agents on alternating blocks of all trial types until each trial type had been seen 5 times. To test predictive inference, we then trained the agents on 500 random trials. For the joint inference algorithm, we used the joint prior and joint task estimate for all of the block trials during training, but not during pretraining or random trials.

For the structural discrimination task, we pretrained agents alternating between blocks of the rewarded trial types until each trial type had been seen 3 times, with each block consisting of 50 trials. We then trained the agents on alternating blocks of all trial types until each trial type had been seen 3 times. To test predictive inference, we then trained the agents on 500 random trials. For the joint inference algorithm, we used the joint prior and joint task estimate for all of the block trials during training, but not during pretraining or random trials.

#### Reinforcement learning problem statement

Reinforcement learning agents aim to maximise reward by interacting with an unknown environment. Here, they aim to solve the randomised cued T-maze, which represents a partially observable Markov-decision process (POMDP) defined by *D* = (*L, A, p, R*, Φ, *γ*) where *L* is the hidden state space, *A* is the action space and Φ is the observation space. For *l* ∈ *L* and *a* ∈ *A* the next hidden state *l*^′^ is given by *p*(*l*^′^| *l, a*). The hidden state *l* is defined by the history of past observations for *ϕ* ∈ Φ. The reward is defined as *R*(*l, a, l*^′^) and *γ* ∈ [0, 1] is the discount factor which reduces reward values with distance into the future. The aim of the agent is to find a policy *π*, a set of mappings from hidden states to actions, that maximises expected discounted total future reward, known as value *V*, based on reward *R*(*s*):

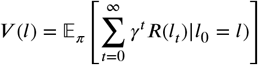

The cued T-maze represents a specific instance of this problem where hidden states factorise into partially observable contexts *Z* and fully observable states *S* which maintain the same transition dynamics *p*(*s*^′^| *s, a*) for all *s* ∈ *S*. The problem then reduces to a set of Markov-decision process (MDPs) *G*^*z*^ = (*S, A, p, R, γ*) for *z* ∈ *Z*, where each context is defined by a specific instantiation of the reward function *r*(*s, a, s*^′^) ∈ *R*(*s, a, s*^′^).

The optimal way to solve this problem is to infer z to predict the current relevant behaviour. However, if *z* cannot be predictively inferred based off of *ϕ* the problem reduces to a transfer learning problem as defined in [41] and [42] where the optimal behaviour is reactive to changes in *R*(*s, a, s*^′^) [43]. To model the behaviour associated with these two distinct strategies, we use an existing hippocampal model that attempts to predict the context and associated best behaviour, and an existing prefrontal cortex model that is reactive to behavioural outcomes.

#### Feature inference model

To estimate value in this set of MDPs, first, the current relevant context must be identified. To infer *z* we implemented an existing model of the hippocampus from [14] (**Figure 8A**). This model estimates its belief about the current context in the form of a belief state *b*, which is a distribution of probabilities over all possible contexts *b*(*z*) = *P* (*z* |*ϕ*). To do so, this model learns a set of trial-type specific successor feature maps *M*^*z*^ where *M* ∈ *M*^*z*^ is a predictive map of how observed features in *ϕ* are related to each other. Concurrently to learning these maps the model uses Bayesian inference over *M*^*z*^ to determine *b*(*z*) based on differing features between contexts.

**Figure 8.**
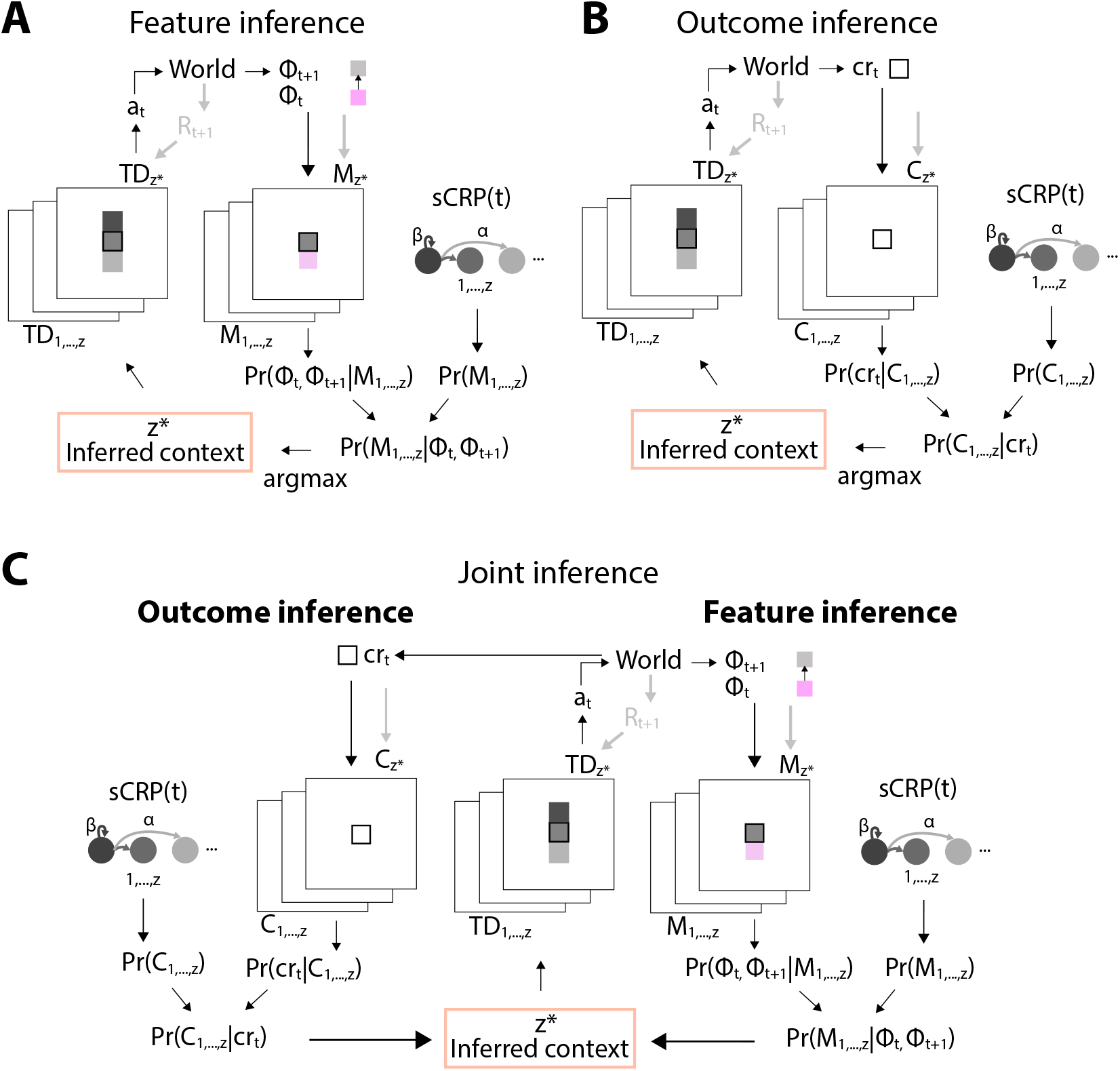
Schematics of model designs. (A) Schematic of HPC model, showing how observed feature transitions are compared with existing successor feature maps using Bayesian inference to determine which context is currently most likely. Schematic of PFC model, (B) showing how observed convolved reward is compared with existing maps of convolved reward using Bayesian inference to determine which context is currently most likely, (C) Schematic of joint model architecture, showing how the context likelihoods generated by both HPC and PFC models are combined for contextual inference and action selection.

Each successor feature map *M* learns the predicted future occupancy of all other features based on the current column vector of observed features *ϕ. M* is an *ixj* matrix that linearly transforms the current feature observations to predict the future occupancy of all features given the current features, called successor features *ψ* (*s*) = *Mϕ*(*s*) for *ϕ* ∈ Φ and *ψ* ∈ Ψ. We use subscript *i* to denote the rows and *j* to denote the columns, such that *M*_*i,j*_ = *M* [*i, j*]. Intuitively, *M*_*i,j*_ is the predicted future occupancy of feature *i* given feature *j. M* is learnt using a simple delta rule *M*_*t*+1_ = *M*_*t*_ + *αδg* after every observation for *t* ∈ *T* with the prediction error:

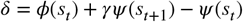

Here, *g*_*i*_ = *ϕ*_*i,t*_/*x*_*t*_ normalises the update depending on feature overrepresentation and the number of features observed x such that entries in *M*_:,*j*_ represent predicted future occupancies for *ϕ*_*j*_ = 1.

From a probabilistic perspective the values *M*_*i,j*_ are point estimates of the future predicted occupancies. We assume their distribution is Gaussian with variance 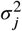, giving a multivariate normal of the form 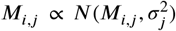 This allows us to use a flexible learning rate known as the Kalman gain *α* = *k*_*t*_ to anneal the learning rate as confidence in *M* increases. The Kalman gain is calculated using a Kalman filter that tracks the uncertainty in model estimates Σ_*t*+1_ = Σ_*t*_ − *k*_*t*_*h*_*t*_Σ_*t*_ versus the uncertainty in observations defined by the observation noise 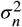:

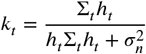

where *h*_*t*_ = *γϕ*_*t*+1_ − *ϕ*_*t*_ is the discounted feature derivative. The Kalman gain decreases as the confidence in feature predictions increases, which is proportional to how often they have been observed.

To find the context map *z* with the best fit to the observed features, Bayesian inference is used: *p*(*z* |*ϕ*_*t*_) ∝ *p*(*ϕ*_*t*_ |*z*)*p*(*z, t*). The map that can best predict *ϕ* is the map where the future predicted occupancies of the observed state transition are best aligned, and thereby is the map that has the smallest prediction error. If *δ* = 0, *ϕ*_*t*_ = *ψ* (*s*_*t*_) − *γψ* (*s*_*t*+1_), so the probability of observing *ϕ* for each context map is estimated with:

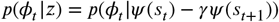

For inference in the feature-based algorithm we assume the variance is a constant predefined hyperparameter 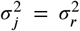 to limit the impact of exploratory actions on context inference, as it allows for an improved tolerance for deviations from the current context map after learning.

For the prior a sticky Chinese restaurant process (sCRP) prior is used. Based on *N*_*z*_, the number of previous observations assigned to each context *z* over the last *y* timesteps, a new context identity is generated according to the sCRP. The sCRP gives the probability of being in any of the previously experienced contexts or being in a new context:

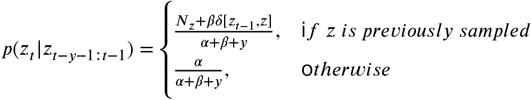

with the concentration *α*, the prior belief as to how likely context switching is, and stickiness *β*, the prior belief as to how likely the context is to stay the same as on the previous step. The sCRP is based on the belief that although there is an infinite number of possible contexts, the agent is most likely to be in the context it has seen the most in the past weighted by recency. Since the prior is intractable, a particle filtering method from [42] is used for its estimation.

The inferred context is thus based on how well the features observed fit the features predicted by the learnt feature map for each context and is given by *z*^∗^ = argmax *p*(*z*|*ϕ*_*t*_).

#### Outcome inference model

As described above, if *M*^*z*^ has not been learnt well or cannot be used to infer the current context the problem nears a transfer learning problem, as inference based on *ϕ* is poor. We used an existing model of PFC from [42] that was designed to solve transfer learning problems to simulate this strategy (**Figure 8B**). The PFC model uses the same framework as the HPC model described above: it learns distinct maps of different contexts and uses Bayesian inference over them to determine the current context, making use of Kalman filtering to have a flexible learning rate and estimate the variance during Bayesian inference. The difference is in what information the maps contain. Instead of learning about observed features of the environment, the PFC model utilises a map of diffuse state-values to separate contexts where rewards are in sufficiently different states, meaning that different policies are necessary to reach rewards. The PFC model estimates using Bayesian inference over a set of convolved reward value maps *C* ∈ *C*^*z*^ that learn a local *b*(*z*) = *P* (*z*|*cr*) value estimate *cr* for each *s* based on reward locations. The PFC model learns about this environment as a tabular environment, where each state can be represented by a one-hot vector defined by current location. Briefly, outcome-based agents use a kernel of the form *F*_*γ*_ = [*γ*^−*len*^, …, *γ*^*len*^] with a filter length *len* either side of the current state to calculate the convolved reward estimate *cr*_*t*_ = *F*_*γ*_ *r*_*t*−2*len*+1:*t*_. Due to the filter, there is a temporal delay of len states preceding inference. The agent updates its predictions for the currently inferred context map using a simple delta rule *C*(*s*)_*t*+1_ = *C*(*s*)_*t*_ + *α*(*cr*_*t*_ − *C*(*s*)_*t*_), where *α* = *α*_*o*_*k*(*s*) and *k* is calculated as above replacing *h*_*t*_ with the current state *s*_*t*_. The estimates the model learns again assume a Gaussian of the form *C*(*s*) ∝ *N* (*C*(*s*), *σ*^2^(*s*)). The same Bayesian inference mechanism is used as in feature inference and *p*(*z* |*cr*_*t*_) is determined with *p*(*cr*_*t*_ |*z*) = *p*(*cr*_*t*_ |*C*(*s*)). As a result, outcome maps have diffuse state value estimates allowing for clustering of different contexts based on different optimal policies.

#### Choice via temporal difference learning

We decided to separate the choice component from the inference component, to compare the ability of feature and outcome inference algorithms specifically in inference. For estimating state-action values and making choices we learn a set of temporal difference maps *T D*^*z*^. Once a context identity has been selected a corresponding temporal difference model is used to pick the optimal action. The temporal difference TD value for each state gives its discounted expected future reward and is updated with *δ*_*t*_ = *r*_*t*+1_ − *T D*_*t*_*ϕ*_*t*_ + *γT D*_*t*_*ϕ*_*t*+1_ and the Kalman gain. The Kalman filter is used as above replacing *h*_*t*_ with the current state features *ϕ*_*t*_. Future predicted reward is given by *Q*(*a*| *ϕ*_*t*_) = *T D*(*ϕ*_*t*+1_) where *ϕ*_*t*+1_ is determined using a one-step lookahead under the assumption that the agent has access to the transition dynamics of the environment *p*(*ϕ*_*t*+1_ |*phi*_*t*_, *a*). The action taken is chosen at random *ϵ* proportion of the time to allow for continuous exploration and with *a*^∗^ = argmax *Q*(*a*|*ϕ*_*t*_) the remainder of the time.

#### Joint inference

During learning, the joint inference algorithm determines the current most likely task based on how well the observations *o*_*t*_ fit both the successor feature maps and convolved reward value maps learnt for each context:

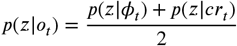

Additionally, a joint prior is used where at each step the prior evolves once according to the feature dynamics and once according to the outcome dynamics. In contrast to maintaining a separate prior for each algorithm, a joint prior aids in ensuring that task estimates converge onto the equivalent maps within each algorithm and allows balancing of the different optimal hyperparameters across different algorithms. The joint inferred context is then used for making choices and updating corresponding feature and outcome maps (**Figure 8C**). After learning, the agent switches to using only feature inference.

#### Other algorithms used for comparison

Here we provide a brief description of the models used for comparison.

#### SR and SR1

The SR algorithms learn one successor representation map M, in the same way as the HPC algorithm does. In contrast to above, we do not learn a separate TD map, but instead learn a reward vector for the SR to make choices. The SR allows decomposition of reward into the reward associated with each feature w and the state features *R*(*s*) = *ϕ*(*s*)*w*. The value function decomposes into future predicted occupancy of all features given the current features and the reward associated with each feature *V* (*s*) = *Mϕ*(*s*)*w. w* is learnt using a simple delta rule with learning rate *α*_*r*_. For one-shot SR *α*_*r*_ is set to 1.

#### TD

The TD algorithm learns one temporal difference map TD as described above.

### Other ways to combine algorithms explored

#### Replay

During each attempt on a trial, the agent keeps track of the feature-transitions it observes. If the agent receives a reward at the end of the attempt, the task identity inferred by the outcome-based model is accepted as ground truth and the feature-transitions are replayed within said task in the feature-based model.

#### Ideal observer

In the ideal observer model the outcome inference algorithm is told the actual task identity, which is assigned a probability of 95 percent. The identity of the other task is assigned a probability of 5 percent.

#### Exploration

Each agent takes 10,000 steps via random walk in the environment without the predictive cues and rewards. The learnt SR and associated covariance make up the random-policy map.

#### Parameters and hyperparameters

Outcome inference and feature inference model parameters were optimised using the initial task with the short maze stem, starting from the original parameters in [42] with an initial manual search for a reasonable range of values followed by a grid search within the found range. Parameters remained the same for all other experiments, apart from the structural discrimination task. As talked about in the discussion, parameters for structural discrimination task were readjusted following the same procedure as above, as there were 6 total contexts instead of 4.

### Statistics

Values plotted are the mean ±95% confidence interval, unless there are outliers that skew the mean in which case the median is plotted (only in Figure 5b: cue-choice distance). Time-series data across algorithms is analysed using mixed ANOVA and bargraph data is analysed using ANOVA. For post-hoc testing, for ANOVAs we used pairwise Tukey tests and for mixed ANOVAs we used multiple T-tests with the Benjamini/Hochberg FDR correction. * indicates p<0.05. Statistical results are detailed in Table 6.

**Table 1.**
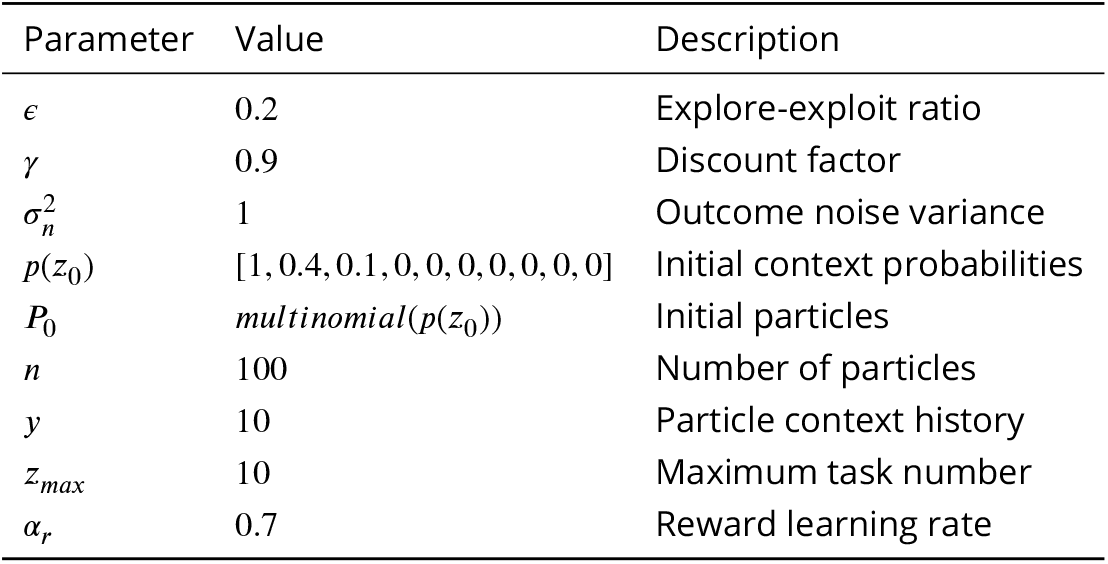
Parameters across all models.

**Table 2.** Parameters for feature inference.

| Parameter | Value | Description |
| --- | --- | --- |
| $\alpha$ | 0.5 | Concentration |
| $\beta$ | 5 | Stickiness |
| $\sigma_0^2$ | 1.6 | Initial residual variance |
| $\sigma_r^2$ | 1.6 | Residual variance for inference |

**Table 3.** Parameters for feature inference in biconditional discrimination.

| Parameter | Value | Description |
| --- | --- | --- |
| $\alpha$ | 0.5 | Concentration |
| $\beta$ | 8 | Stickiness |
| $\sigma_0^2$ | 1.6 | Initial residual variance |
| $\sigma_r^2$ | 1.6 | Residual variance for inference |

**Table 4.** Parameters for outcome inference.

| Parameter | Value | Description |
| --- | --- | --- |
| $\alpha$ | 0.1 | Concentration |
| $\beta$ | 0.7 | Stickiness |
| $\sigma_0^2$ | 1.6 | Initial residual variance |
| $\alpha_o$ | $\frac{2}{3}$ | Learning rate weighting |
| $len$ | 2 | Filter length |

**Table 5.** Parameters for outcome inference in biconditional discrimination.

| Parameter | Value | Description |
| --- | --- | --- |
| $\alpha$ | 0.8 | Concentration |
| $\beta$ | 0.7 | Stickiness |
| $\sigma_0^2$ | 1.6 | Initial residual variance |
| $\alpha_o$ | $\frac{2}{3}$ | Learning rate weighting |
| $len$ | 2 | Filter length |

**Table 6.**
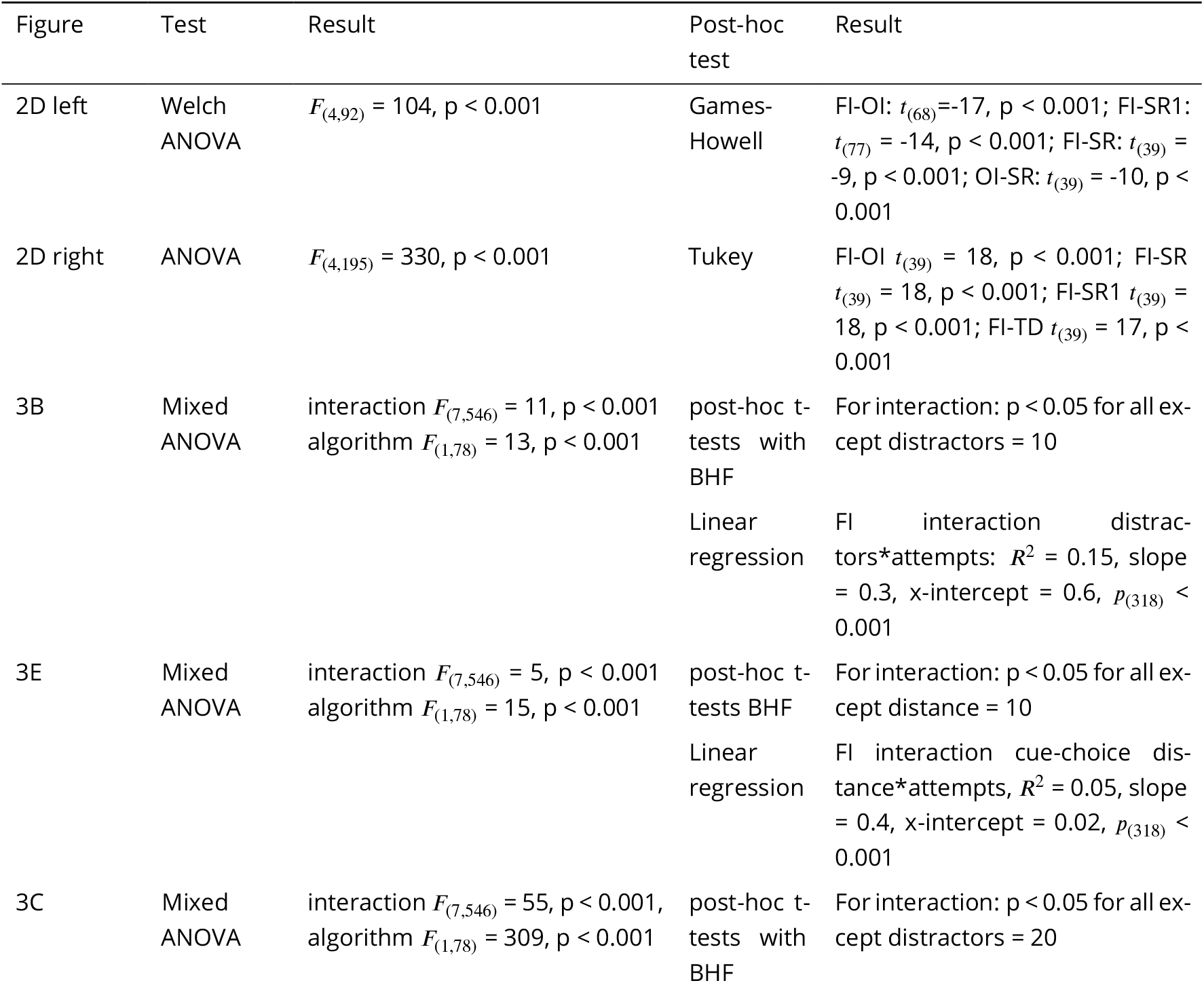
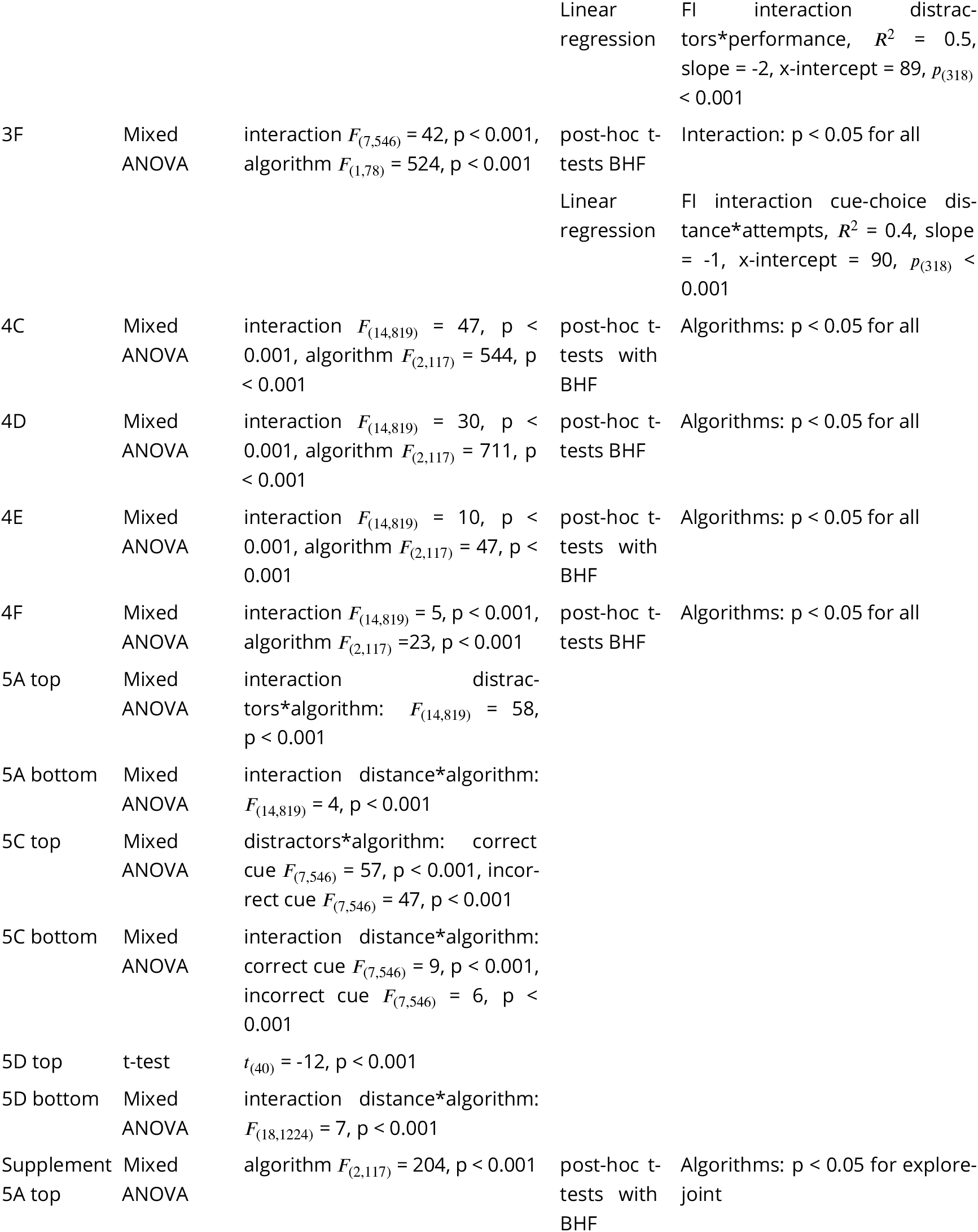
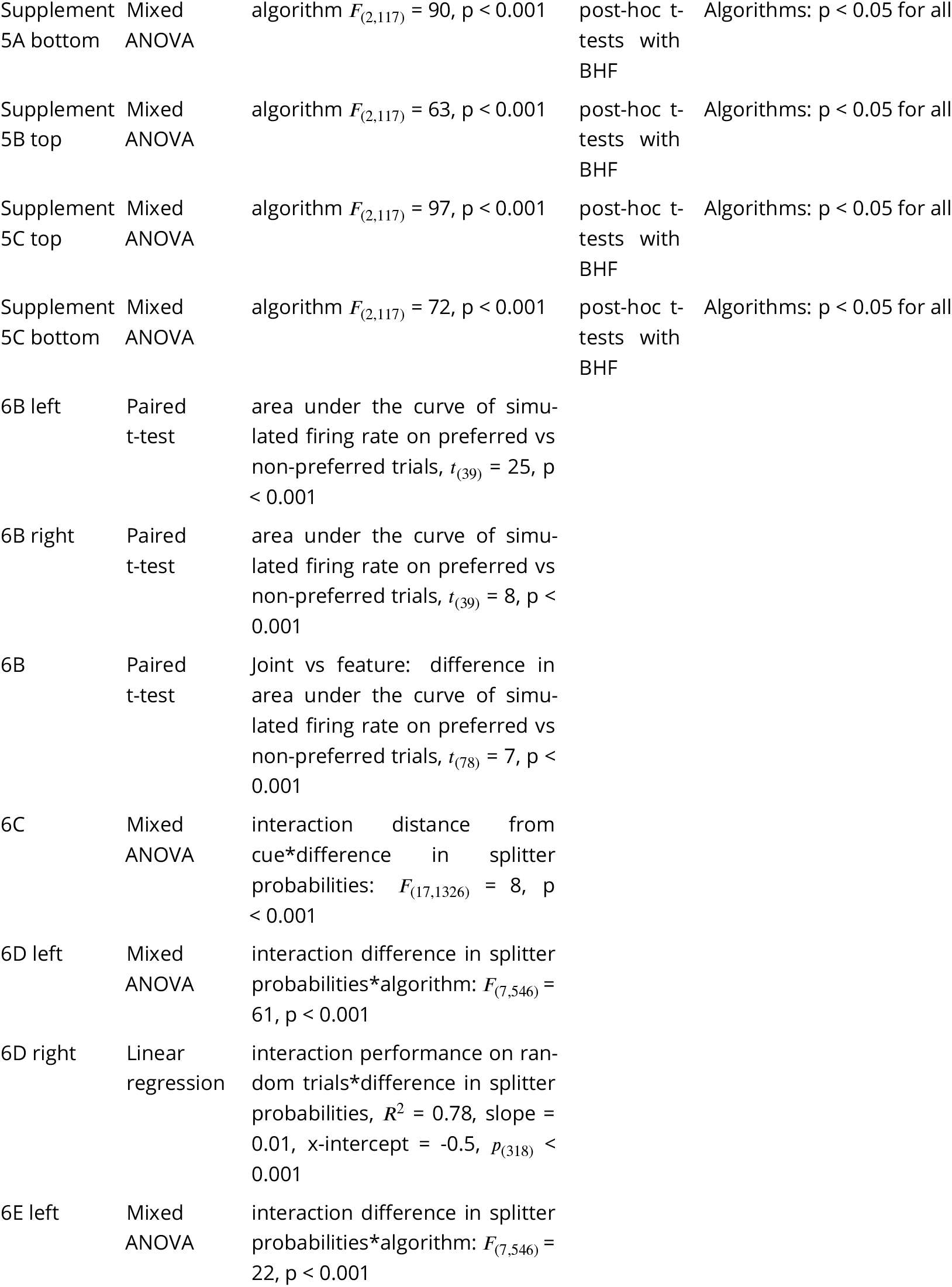
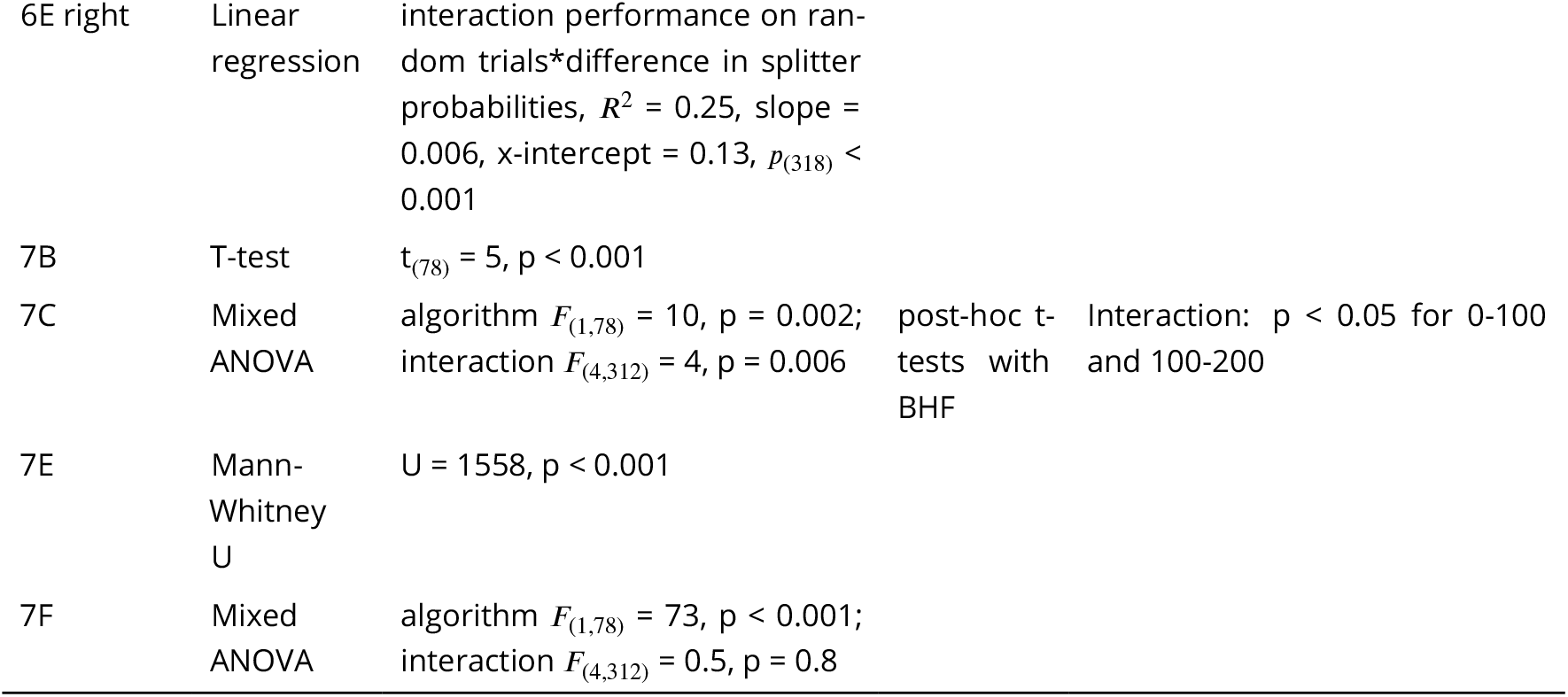
Statistical results.

### Compute power

The compute power needed to generate the data in this paper from these algorithms is around 300 hours using 200 cores each with 5GB RAM.

### Code and data availability

Model code, analysis code and data generated by the models and used for analysis will be made available publicly.

## ACKNOWLEDGMENTS

We thank Daniel Bush, Jesse Geerts, Neil Burgess and other members of the MacAskill and Burgess labs for helpful discussions and feedback. A.F.M. was supported by a Sir Henry Dale Fellowship jointly funded by the Wellcome Trust and the Royal Society 109360/Z/15/Z; and by an MRC project grant MR/W02005X/1. J.P. was supported by the Wellcome Trust 4-year PhD in Neuroscience at UCL 222292/Z/20/Z.

---

**Figure 4—figure supplement 1.**
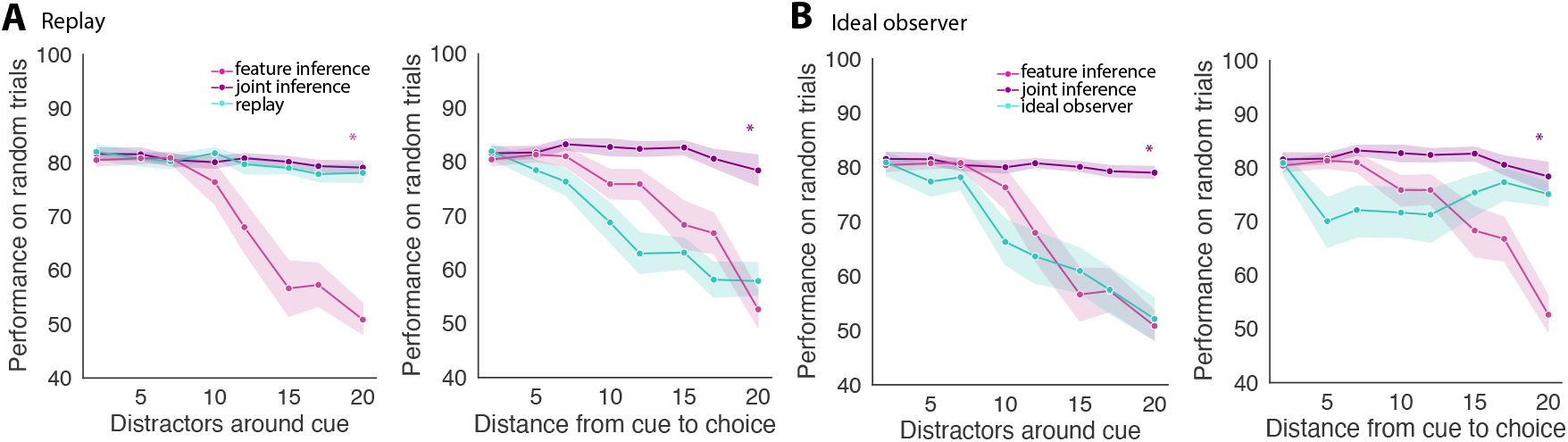
Inconsistent performance of algorithms using other ways of combining outcome inference and feature inference during learning. Performance on random trials with increasing distractors around cue or increasing distance from cue to choice using (A) replay of trial observations into the context map inferred by outcome inference at the end of trials, (B) replacing the outcome inference algorithm with an ideal observer model.

**Figure 5—figure supplement 1.**
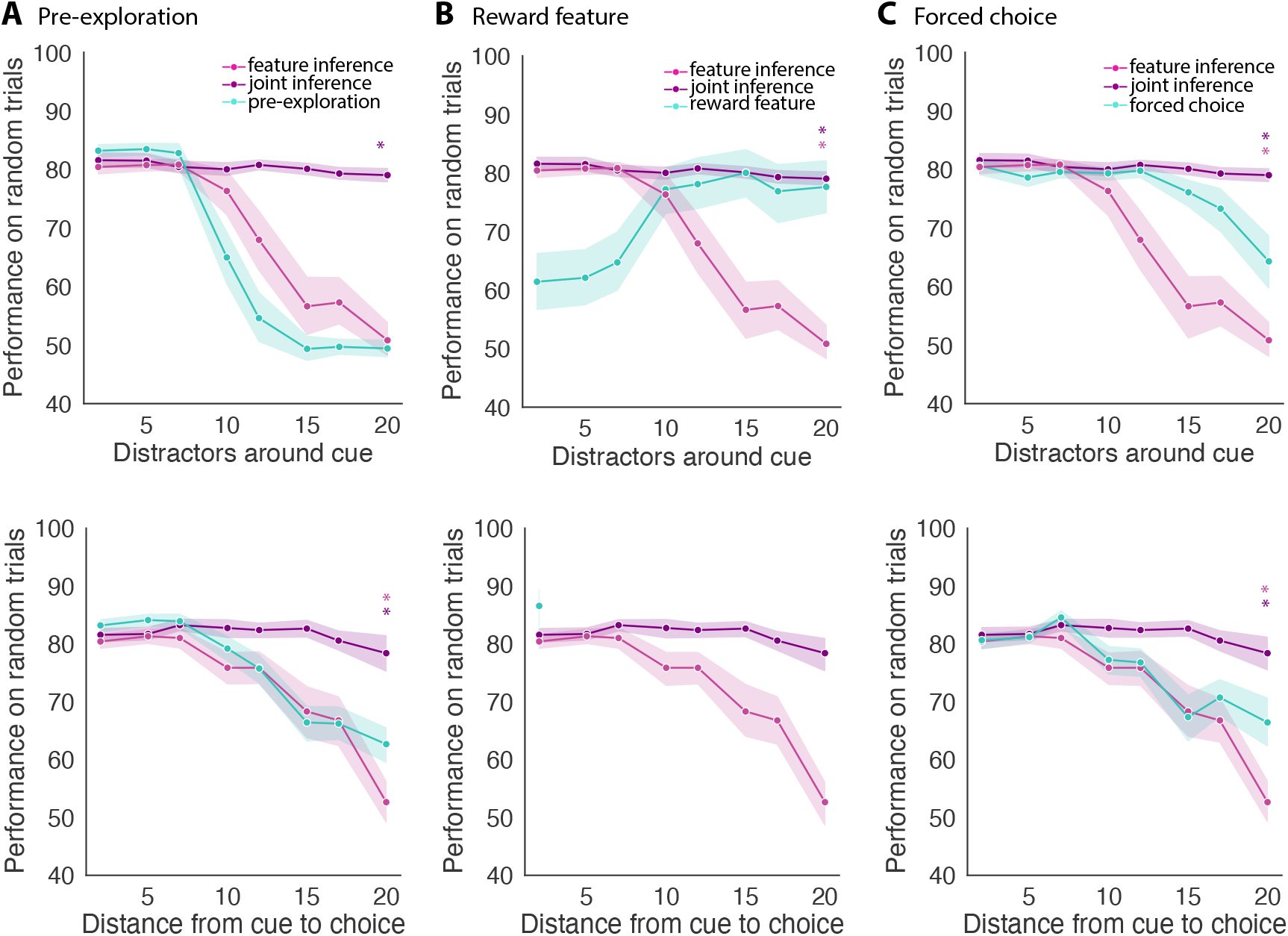
Inconsistent performance of algorithms using methods other than outcome-inference to support learning. Performance on random trials with increasing distractors around cue or increasing distance from cue to choice using (A) pre-exploration of the environment without rewards or predictive cues, (B) an additional feature that indicates reward presence (the data in cue-choice distance is incomplete as individual agents took over the 48hr time limit to run), (C) forced choice where the incorrect arm is blocked off during learning.

## Notes

### Competing Interest Statement

The authors have declared no competing interest.

